# Phosphorylation of the AP2 μ2 subunit by p70S6 kinase facilitates clathrin-mediated endocytosis

**DOI:** 10.1101/2025.02.13.638021

**Authors:** Aleksandra Tempes, Agnieszka Brzozowska, Tomasz Węgierski, Ayomide Fasemire, Katarzyna Olek, Kamil Jastrzębski, Ewa Liszewska, Katarzyna Misztal, Katarzyna Machnicka, Matylda Macias, Aleksandra Szybińska, Ewa Sitkiewicz, Agata Malinowska, Agata Gozdz, Agnieszka Wyszyńska, Anna Łasica, Marta Hoffmann-Młodzianowska, Katarzyna Orzoł, Marta Miączyńska, Maria W Górna, Wojciech Pokrzywa, Jacek Jaworski, Anna R Malik

## Abstract

Clathrin-mediated endocytosis (CME) internalizes cell-surface receptors via clathrin-coated invaginations of the plasma membrane. Both clathrin and endocytic cargo are recruited to these sites by the adaptor protein complex AP2. AP2 cycles between a closed cytoplasmic conformation and an open membrane-bound state, and efficient CME requires both conformations and their dynamic interconversion. The mechanisms regulating these conformational changes, which include post-translational modifications of the AP2, remain incompletely understood. Here, we report that p70S6 kinase phosphorylates the µ2 subunit of the AP2 and that the phosphorylation of serine 45 (S45) depends on p70S6K activity. Loss of S45-µ2 phosphorylation results in decreased internalization of canonical CME cargo such as transferrin and PDGF receptors. In *Caenorhabditis elegans*, lack of S45-µ2 phosphorylation produces directionally similar but markedly weaker phenotypes than AP2 loss of function. Live imaging and *in silico* dynamic modelling suggest that S45-μ2 phosphorylation has impact on the conformational changes of the AP2 complex. These findings identify a p70S6K-dependent mechanism that modulates AP2 function and further strengthen the importance of post-translational regulation in controlling CME.

**Summary statement:** Clathrin-mediated endocytosis is essential for receptor internalization and cell signaling. This study identifies a novel regulatory mechanism in which p70S6K-mediated phosphorylation of the AP2 subunit μ2 at Ser45 modulates AP2 conformational dynamics and facilitates receptor endocytosis.

## Introduction

The intracellular environment of mammalian cells is protected from the extracellular environment by the plasma membrane, which is impermeable to hydrophilic molecules. Because of this strict protection, specialized mechanisms such as endocytosis need to be utilized for the uptake of molecules critical for proper cell function. Among various types of endocytosis, clathrin-mediated endocytosis (CME) is the most intensively studied due to its key role in development, physiology, and its involvement in diseases such as neurodegeneration and cancer (Collins *et al*, 2002; Jackson *et al*, 2010). During CME, cargo is internalized from the plasma membrane via so-called clathrin-coated pits (CCPs) either constitutively (e.g., transferrin receptor, TrfR) or exclusively upon ligand binding (e.g., platelet-derived growth factor receptor, PDGFR). CCPs are invaginations of the plasma membrane enriched with the cargo destined for internalization. They are covered with a clathrin lattice, which shapes CCPs into clathrin-coated vesicles (CCVs) (Kadlecova *et al*, 2017).

However, clathrin on its own cannot bind either cargo or the plasma membrane. Therefore, for CME to take place, the adaptor protein complex AP2 is utilized, which simultaneously binds to the plasma membrane, cargo, and clathrin (Collins *et al*, 2002; Jackson *et al*, 2010). AP2 consists of four subunits: two large ones (α and β2 [∼ 110 kDa]), medium µ2 (50 kDa), and small σ2 (17 kDa). While all four constitute the core of AP2, the α and β2 subunits extend the hinge and appendage domains (or α-and β2-ears) beyond the core. The α, β2, and µ2 subunits recognize phosphatidylinositol 4,5-bisphosphate (PIP2) and are therefore responsible for AP2 binding to the plasma membrane (Jackson *et al*, 2010; Kadlecova *et al*, 2017). Finally, the µ2 and σ2 acting together with α subunit bind cargo possessing Yxxφ and dileucine motifs, respectively. The AP2 core can adopt several different conformations including closed and open conformation. In the closed conformation, µ2 and β2 domains, responsible for binding cargo and clathrin, respectively, are inaccessible. Upon binding PIP2 at the plasma membrane, AP2 opens to bind cargo and clathrin (Collins *et al*, 2002; Jackson *et al*, 2010; Beacham *et al*, 2018; Kelly *et al*, 2008; Kovtun *et al*, 2020; Wrobel *et al*, 2019).

CME is strictly controlled, and many key endocytic proteins are phosphorylated at some point (Korolchuk & Banting, 2003; Liberali *et al*, 2008). Endocytosis-regulating phosphorylation can affect CME progress either positively or negatively (Korolchuk & Banting, 2003; Liberali *et al*, 2008). Phosphorylation of µ2 at threonine 156 (T156) is the most extensively studied one, but its exact function is still debated (Wrobel *et al*, 2019; Partlow *et al*, 2019; Fingerhut *et al*, 2001; Ricotta *et al*, 2002). Of note, according to PhosphoSitePlus (https://www.phosphosite.org), 16 additional serine or threonine phosphorylation sites were detected in µ2 by mass spectrometry. Of them, only the role of serine 45 (S45) phosphorylation was partially characterized, suggesting its negative role in the internalization of the transient receptor potential cation channel subfamily V member 1 (TRPV1) (Liu *et al*, 2019).

One of the pathways potentially involved in regulating CME is the mammalian target of rapamycin (mTOR) signaling pathway (Malik *et al*, 2013). mTOR signaling plays a central role in integrating the metabolic status of the cell with extracellular cues, thereby coordinating key processes such as protein synthesis, cytoskeletal dynamics, and autophagy. A high-throughput screening study has pointed to a role of mTOR in the regulation of CME (Pelkmans *et al*, 2005). This finding was further supported by observations that nutrient deprivation, a condition that strongly suppresses mTOR activity, impairs CME by arresting the formation of CCPs and reducing transferrin uptake (Colombero *et al*, 2021). Despite these observations, the precise molecular mechanisms connecting mTOR signaling to the regulation of CME remain unclear.

One of the primary effectors of mTOR is p70S6 kinase (p70S6K), which is best known for its role in the regulation of protein synthesis. Its functions are mostly studied in the context of mTORopathies, a group of diseases resulting from excessive activity of the mTOR kinase and its downstream targets, including p70S6K (Karalis & Bateup, 2021; Switon *et al*, 2017). In most cases, mTOR overactivation is caused by inactivating mutations of its key regulators, *TSC1* or *TSC2*, encoding hamartin and tuberin, respectively. Recent phosphoproteomics data have revealed hundreds of proteins phosphorylated in a p70S6K-dependent manner, pointing to its role in the regulation of a variety of cellular processes in addition to translation, including control of protein folding, RNA splicing, nucleotide synthesis, cytoskeleton dynamics, and membrane trafficking (Bonucci *et al*, 2020; Fricke *et al*, 2023; Fumagalli & Pende, 2022). Here, we show that p70S6K may be a key mediator of mTOR’s impact on CME. Using various cargo-uptake assays, live imaging of CCP formation, and mass spectrometry, we investigated the role of p70S6K in CME regulation. We showed that p70S6K activity is required for CME cargo uptake in human cells, an effect mediated by p70S6K-dependent phosphorylation of µ2 at S45. This modification of µ2 is likely essential for AP2 conformational changes and the efficient formation of CCPs.

## Results

### Cargo internalization via CME is upregulated in cells with increased p70S6K activity

Evidence exists for a p70S6K contribution to CCV formation in very specialized cells (proximal tubular cells) (Grahammer *et al*, 2017), but it is unclear if and how p70S6K affects this type of endocytosis in other cell types. Therefore, we first looked at CME in cells with increased p70S6K activity due to loss of hamartin (for review see: (Switon *et al*, 2017; Malik *et al*, 2013; Switon *et al*, 2016)). First, we generated HeLa cell lines stably expressing shRNA targeting *TSC1* mRNA (HeLa^shTSC1^) and GFP-tagged clathrin light chain (EGFP-CLCa) to enable further detailed analysis of CCPs formation. As a control, we generated HeLa^WT^ cell line expressing EGFP-CLCa. Constitutive overexpression of shTSC1 resulted in an almost complete loss of hamartin (**Fig. 1A, B**) and, as anticipated, an increase in phosphorylation of the direct target of p70S6K, ribosomal protein S6 (P-S6, Ser235/236), confirming the activation of p70S6K (**Fig. S1A, B**). Next, we tested whether the knockdown of *TSC1* affects the early steps of the CME by analyzing the uptake of fluorescently labeled transferrin. Transferrin uptake occurs via the transferrin receptor (TrfR), a canonical CME cargo. Cells with decreased *TSC1* expression and overactivation of p70S6K showed an increased uptake of transferrin compared to control cells (**Fig. 1C, D**). Pretreatment of the cells with a p70S6K inhibitor (p70S6Ki) clearly prevented the increased transferrin uptake induced by *TSC1* knockdown (**Fig. 1C, D**), substantiating the notion that the activity of p70S6K is critical for enhanced CME seen in HeLa^shTSC1^.

**Figure 1.**
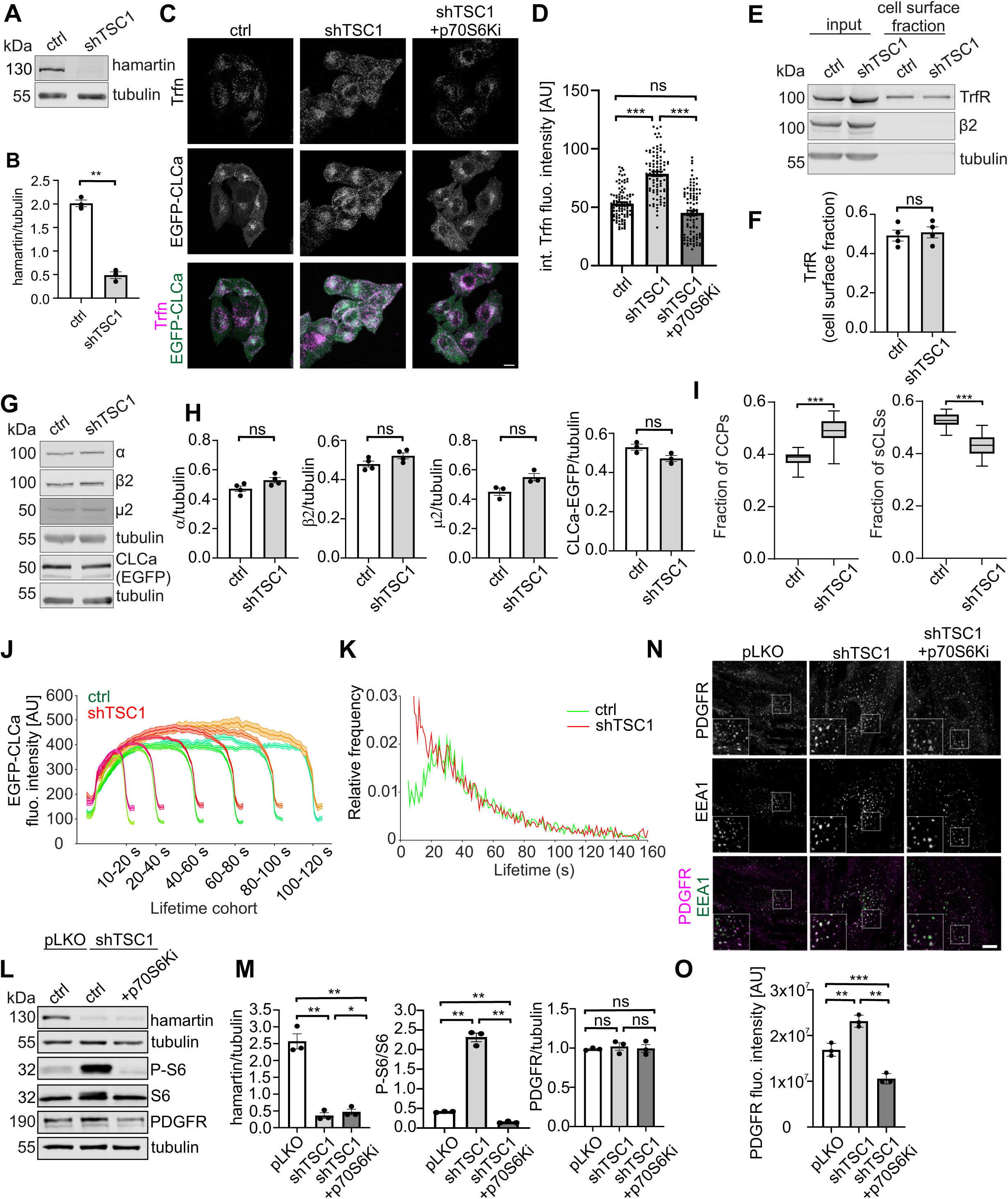
Cargo internalization via CME is upregulated in cells with overactive p70S6K. **A.** Western blot showing levels of endogenous hamartin in HeLa cells stably expressing EGFP-CLCa only (ctrl) or with TSC1 shRNA (shTSC1). **B.** Quantitative analysis of hamartin levels. *N* = 3 independent experiments. \*\**p* < 0.01 (paired t-test; t = 10.81, df = 2). **C.** Representative confocal images of cells after 5-minute uptake of Alexa Fluor 647-labeled transferrin (Trfn, magenta). HeLa cells stably expressing EGFP-CLCa only (ctrl) or together with TSC1 shRNA (shTSC1) were serum-starved and treated with 0.1% DMSO or 10 µM p70S6K inhibitor for 2 h before transferrin uptake. EGFP-CLCa is shown in green. Scale bar = 10 µm. **D.** Analysis of immunofluorescence signal of internalized Alexa Fluor 647-conjugated transferrin normalized to cell area in cells treated as in *C*. The data are shown as the mean value ± SEM. *N* = 2 independent experiments. Number of cells (*n*) = 101 (ctrl), 93 (shTSC1), 96 (shTSC1 +p70S6Ki). \*\*\**p* < 0.001, *ns* – non-significant (Kruskal Wallis test [H = 107.3] with Dunn’s multiple comparisons test). **E.** Western blot showing cell membrane levels of TrfR in HeLa cells as in *A*, analyzed by pull-down of biotinylated cell surface proteins. **F.** Quantitative analysis of TrfR surface expression. *N* = 4 independent experiments. *ns* – non-significant (paired t-test; [t = 0.2918, df = 3]). **G.** Western blot showing expression level of AP2 subunits and clathrin light chain fused with EGFP (EGFP-CLCa) in cell lines as in *A*. **H.** Quantitative analysis of AP2 α, β2, µ2 subunits, and CLCa-EGFP levels. *N* = 4 independent experiments for α, β2. *N* = 3 independent experiments for µ2 and CLCa-EGFP. *ns* – non-significant (paired t-test; α [t = 1.587, df = 3], β2 [t = 1.376, df = 3], µ2 [t = 1.994, df = 2], CLCa-EGFP [t = 1.848, df = 2]). **I.** Fractions of all detected CCPs and sCLSs found among all valid tracks (ctrl: number of analyzed movies (*n*) = 26, number of total valid tracks = 27800; shTSC1: number of analyzed movies (*n*) = 30, number of total valid tracks = 25875). Box plots show medians, 25th and 75th percentiles, and outermost data points. \*\*\**p* < 0.001 Matlab ranksum (*p* value 1.92e-09 for CCPs and 5.58e-10 for sCLSs). **J.** Clathrin fluorescence intensity in CCPs lifetime cohorts of control cells (green) and cells with *TSC1* knockdown (red). Intensities are shown as mean ± SE. **K.** Lifetime distributions of all CCPs found in HeLa control cells (green) and with *TSC1* knockdown (red). **L.** Western blot showing levels of endogenous hamartin, P-S6 (235/236), S6, PDGFR and tubulin in CCD-1070Sk cells transduced with control (pLKO) or shTSC1-encoding lentiviral vector, serum-starved and treated with 0.1% DMSO or 10 µM p70S6K inhibitor for 2 h. **M.** Quantitative analysis of hamartin, P-S6 (S235/236) and PDGFR levels. *N* = 3 independent experiments. \*\**p* < 0.01, \**p* < 0.05, *ns* – non-significant (RM one way ANOVA followed by Tukey’s multiple comparisons test). **N.** Representative confocal images of CCD-1070Sk treated as in *L*, stimulated with PDGF for 15 minutes and immunofluorescently stained for PDGFR (magenta) and EEA1 (green). Scale bar = 10 µm. **O.** Analysis of PDGFR endocytic immunofluorescence signal intensity in cells treated as in *N*. The data are shown as the mean value ± SEM. *N* = 3 independent experiments. \*\*\**p* < 0.001, \*\**p* < 0.01 (One-way ANOVA followed by Tukey’s multiple comparisons test, pLKO vs. shTSC1: q = 8.367, df = 6; pLKO vs. shTSC1+p70S6Ki: q= 8.451, df = 6; shTSC1 vs. shTSC1+p70S6Ki: q = 16.82, df = 6).

Next, we looked for the causes of the difference in transferrin uptake between HeLa^WT^ and HeLa^shTSC1^ cells. First, we investigated whether changed levels of surface TrfR could explain the increased transferrin uptake. Western blot analysis of membrane protein fraction obtained from HeLa^WT^ and HeLa^shTSC1^ cells using the cell surface biotinylation assay revealed no significant reduction of TrfR cell surface levels (**Fig. 1E, F**). The silencing of *TSC1* also did not affect the amounts of AP2 complex subunits (**Fig. 1G, H**). Therefore, we tested whether the increased internalization of transferrin is associated with altered dynamics of early CME steps in cells with TSC1 depletion and p70S6K overactivation. To test this hypothesis, we performed an analysis of CCV formation using total internal reflection fluorescence (TIRF) microscopy based on the behavior of the GFP-tagged clathrin light chain (EGFP-CLCa). TIRF microscopy allows visualization of clathrin-coated pits (CCPs) at the cell membrane and measurement of the parameters describing all steps of CCP formation: nucleation, initiation, growth, and maturation (Loerke *et al*, 2009; Merrifield *et al*, 2002; Mettlen & Danuser, 2014). More specifically, we analyzed the number of subthreshold clathrin-labeled structures (sCLSs) and CCPs, the rate of clathrin polymerization, and finally the lifetime of CCPs. sCLSs s are short-lived (up to 15 s) and dim structures, which never reach maturation and in consequence are aborted. In contrast, mean lifetime of CCPs is >20 s to even 2-3 minutes and their fluorescence, reflecting clathrin recruitment, increases during maturation and then decreases due to scission from the plasma membrane (Aguet *et al*, 2013; Loerke *et al*, 2011). A high number of sCLSs exceeding the number of CCPs may be a sign that CCP initiation is frequent but fails to proceed to the mature state, for example due to improper recruitment of adaptor proteins. In contrast, a high number of CCPs relative to total clathrin structures may indicate productive and efficient endocytosis. The rate of clathrin polymerization, measured as the slope of EGFP-CLCa fluorescence intensity curve at the initial growth phase, provides information about the speed of clathrin incorporation into nascent CCP. The speed of clathrin incorporation is critical to the entire process of CME: too slow recruitment results in the abortion of the forming CCP (Kadlecova *et al*, 2017). Moreover, the differences in EGFP-CLCa signal intensities at the starting point give information about the initial size of forming CCP (Aguet *et al*, 2013). The results of our analysis indicated important differences in the dynamics of CCPs formation between HeLa^shTSC1^ and control cells. Thus, in HeLa^shTSC1^ cells we observed more bona fide CCPs and fewer sCLSs than in control cells (**Fig. 1I**). Moreover, the starting point for EGFP fluorescence in HeLa^shTSC1^ cells was higher when compared to control cells **(Fig. 1J**), which suggests that the CCPs are bigger. Cells depleted of TSC1 exhibited an increase in the fraction of short-lived CCPs (≤40 s) compared with control cells (**Fig. 1K**). Not all CCPs go through the whole life cycle: even 30-50 % of short-lived CCPs fail to mature (Mettlen *et al*, 2018), but HeLa^shTSC1^ cells internalized transferrin more efficiently than control cells, which suggests that a fraction of those short-lived CCPs is fully functional in these cells. This could also provide an explanation for the higher amount of CCPs in those cells when compared to control cells. Altogether, our TIRF microscopy analysis suggests that in cells with lowered *TSC1* expression and p70S6K overactivation, the whole process of CME is more efficient compared to the cells with normal levels of hamartin.

Knowing that increased p70S6K activity in HeLa^shTSC1^ cells enhances the CME of TrfR, we sought to determine whether this effect extends to another CME cargo and to a different cell type. To this end, we analyzed PDGFR internalization (Jastrzębski *et al*, 2017) in CCD-1070Sk fibroblasts stimulated with its ligand PDGF. As before in HeLa cells, we depleted these cells of hamartin to drive overactivation of the mTOR-p70S6K signaling. Towards this end, we used lentiviral delivery of shTSC1 to generate CCD-1070Sk^shTSC1^ cells while the control cells received the pLKO vector (**Fig. 1L, M)**. As in the transferrin uptake in HeLa cells described above, CCD-1070Sk cells with overactive p70S6K endocytosed significantly more PDGFR than control cells upon PDGF treatment (**Fig. 1N, O)** as revealed by immunofluorescence intensity analysis for internalized PDGFR colocalizing with EEA1, a marker of early endosomes. In non-stimulated cells, we did not note PDGFR-EEA1 colocalization, in line with the model of PDGF-induced receptor internalization (Fig. S1C). At the same time, the total PDGFR level remained unchanged compared with control cells, while P-S6 was increased (**Fig. 1L, M**). Like in the case of TrfR endocytosis in HeLa cells, increased PDGFR internalization in CCD-1070Sk^shTSC1^ cells was abolished by pretreatment with p70S6Ki (**Fig. 1N, O**). Overall, we concluded that p70S6K is responsible for the upregulation of early CME steps in cells with *TSC1* knockdown, both in the case of TrfR internalization in HeLa cells and PDGFR endocytosis in fibroblasts.

### Efficient CME requires p70S6K activity in cells with basal mTOR activity

Our discovery that p70S6K is involved in enhanced cargo uptake via CME in cells with hyperactive mTOR prompted us to check if the same holds true in cells with intact *TSC1*. Towards this end, we performed the transferrin uptake assay in HeLa cells with reduced expression of the µ2 encoding gene *AP2M1* (HeLa KO µ2) (Chen *et al*, 2017) transfected with a plasmid encoding myc-μ2^WT^ (Motley *et al*, 2006) along with EGFP to label the transfected cells and visualize their morphology. An analogous approach, i.e. replacement of endogenous µ2 with exogenous one, was used in the work of Kadlecova *et al*. (2017), as it allows a more accurate comparison of the effects of mutations in µ2 on the progression of CME and a better insight into the stage of impairment of AP2 functions. Obtained μ2^WT^ cells were kept serum-starved before the addition of p70S6K inhibitor or DMSO (negative control), and then transferrin internalization was measured as described above for HeLa^shTSC1^. Treatment with p70S6Ki decreased P-S6 levels compared with control cells, confirming its efficacy **(Fig. 2A, B)**. The amount of internalized transferrin was also lower after treatment with p70S6Ki (**Fig. 2C, D**). To further confirm this finding, we used an alternative cytometry-based approach and monitored transferrin uptake over time following p70S6K inhibition. These experiments confirmed decreased transferrin internalization in cells treated with p70S6Ki. Notably, this effect diminished over time and was observed at 5, but not at the 10-and 15-minute time points (Fig. S2A-B).Reduced transferrin uptake was not due to decreased TrfR levels, as determined by Western blotting and immunofluorescence (**Fig. 2A, B, E, F**). We also tested whether inhibition of p70S6K activity affected the levels of transferrin binding to the cell surface, as an estimate of the TrfR presence. To this end, we treated μ2^WT^ cells with p70S6Ki as described above and assessed the levels of fluorescently labeled transferrin bound to the cell membrane. As a result, we observed a slightly increased transferrin fluorescence on the cell surface in μ2^WT^ cells treated with p70S6Ki. Thus, decreased transferrin internalization in cells treated with p70S6Ki is not an effect of the lower amount of TrfR on the cell surface (**Fig. 2G, H**). p70S6Ki treatment also did not affect the levels of overexpressed myc-μ2^WT^ and the endogenous β2 subunit, ruling out the possibility that the differences in transferrin uptake were due to unequal AP2 levels (**Fig. 2A, B**). Thus, we conclude that the p70S6Ki treatment of HeLa cells results in decreased internalization of TrfR. To extend our observation to other cells and cargoes, we checked whether endocytosis of the PDGFR in wild-type CCD-1070Sk fibroblasts also depends on intact p70S6K activity. Indeed, p70S6Ki treatment resulted in a significant decrease in internalized PDGFR after PDGF stimulation (**Fig. 2I, J**). At the same time, p70S6Ki completely blocked the phosphorylation of S6 protein, while the total PDGFR level barely changed compared with control cells (**Fig. 2K, L**). Thus, we conclude that inhibition of p70S6K impairs CME, and this effect is observed in two different cell types and with two CME cargoes studied.

**Figure 2.**
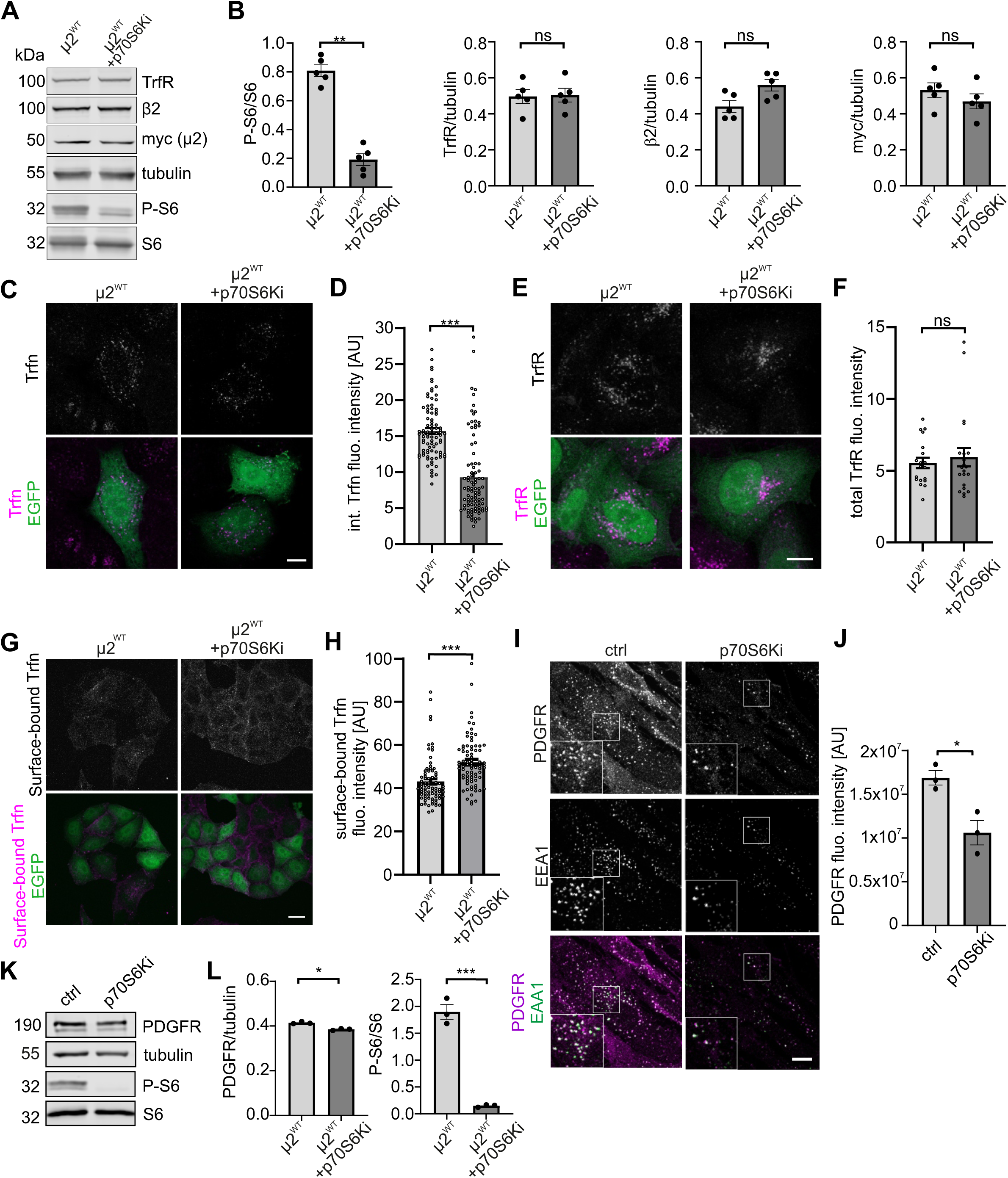
Efficient CME requires p70S6K activity in cells with basal mTOR activity. **A.** Western blot showing levels of endogenous TrfR, β2, S6, P-S6, and tubulin in HeLa cells transfected with plasmids encoding myc-μ2^WT^ and EGFP, and treated with 0.1% DMSO or p70S6K inhibitor (p70S6Ki, 10 μM, 2 h) after overnight starvation in FBS-free medium. **B.** Quantitative analysis of TrfR, β2, myc-µ2, and P-S6 (S235/236) levels. *N* = 5 independent experiments. \*\**p* < 0.01, *ns* – non-significant (paired t-test; TrfR [t = 0.08916, df = 4], β2 [t = 1.842, df = 4], myc-µ2 [t = 0.7428, df = 4], P-S6 [t = 7.540, df = 4]). **C.** Representative confocal images of cells after transferrin uptake. HeLa KO μ2 transfected with plasmids encoding myc-μ2^WT^ and EGFP were treated for 2 h with 0.1% DMSO or 10 µM p70S6Ki before 5-minute incubation with Alexa Fluor 647-labeled transferrin. Transferrin is seen in magenta and EGFP in green. Scale bar = 10 μm. **D.** Quantification of the fluorescence signal intensity of transferrin internalized by cells treated as in *C*. Data are presented as mean ± SEM. *N* = 3 independent experiments. Number of analyzed cells (*n*) = 80 (μ2^WT^), 87 (μ2^WT^ + p70S6Ki), \*\*\**p* < 0.001 (Student’s t test for independent samples; t = 8.728, df = 174). **E.** Representative confocal images of HeLa KO μ2 cells transfected with plasmids encoding myc-μ2^WT^ and EGFP, serum-starved and treated for 2 h with 0.1% DMSO, or 10 µM p70S6Ki. Cells were immunofluorescently stained for transferrin receptor (TrfR, magenta) and EGFP (green). Scale bar = 10 μm. **F.** Quantification of the TrfR fluorescence signal intensity level in cells treated as in *E*. Data are presented as mean ± SEM. *N* = 2 independent experiments. Number of analyzed cells (*n*) = 19 (DMSO), 21 (p70S6Ki). *ns* – non-significant (Mann-Whitney test, U = 195). **G.** Representative confocal images of Alexa Fluor 647-conjugated transferrin (magenta) bound to cell surface of HeLa KO μ2 cells transfected with plasmids encoding myc-μ2^WT^ and EGFP (green), serum starved and treated with 0.1% DMSO or p70S6Ki (10 μM, 2 h). Scale bar = 20 μm. **H.** Quantification of the mean immunofluorescence intensity of transferrin conjugated with Alexa Fluor 647 bound to cell surface. Cells were treated as in *G*. The data are expressed as mean ± SEM. *N* = 4 independent experiments. Number of analyzed cells (*n*) = 78 (μ2^WT^), 81 (μ2^WT^ + p70S6Ki). \*\*\**p* < 0.001 (Mann-Whitney test, U = 1487). **I.** Representative confocal images of CCD-1070Sk cells that were serum-starved, treated with 0.1% DMSO (ctrl) or 10 µM p70S6Ki for 2 h and stimulated with PDGF for 15 minutes. Cells were immunofluorescently stained for PDGFR (magenta) and EEA1 (green). Scale bar = 10 µm. **J.** Analysis of endocytic PDGFR immunofluorescence signal intensity in cells treated as in *I*. The data are shown as the mean value ± SEM. *N* = 3 independent experiments. \**p* < 0.05 (unpaired t-test for, t = 3,894, df = 4). **K.** Western blot showing levels of P-S6 (235/236), S6, PDGFR, and tubulin in CCD-1070Sk cells that were serum-starved and treated with 0.1% DMSO (ctrl) or 10 µM p70S6Ki for 2 h. **L.** Quantitative analysis of PDGFR and P-S6 (S235/236) levels in cells treated as in *K*. *N* = 3 independent experiments. \**p* < 0.05, \*\*\**p* < 0.001 (paired t-test, PDGFR [t = 5.779, df = 2], P-S6 [t = 11.65, df = 2]).

### μ2 is phosphorylated by p70S6K

Given the results described above and the potential regulatory role of μ2 phosphorylation by p70S6K in CME progression, we first checked the ability of the two proteins to interact. Towards this end, we purified GFP-Avi-tag-p70S6K or GFP2-Avi-tag-βGal overexpressed in HEK cells and incubated with recombinant his-tagged μ2 **(Fig. 3A**). We documented interaction of μ2 with GFP-p70S6K while we observed no binding of μ2 to the control resin with immobilized GFP-βGal (**Fig. 3A**).

**Figure 3.**
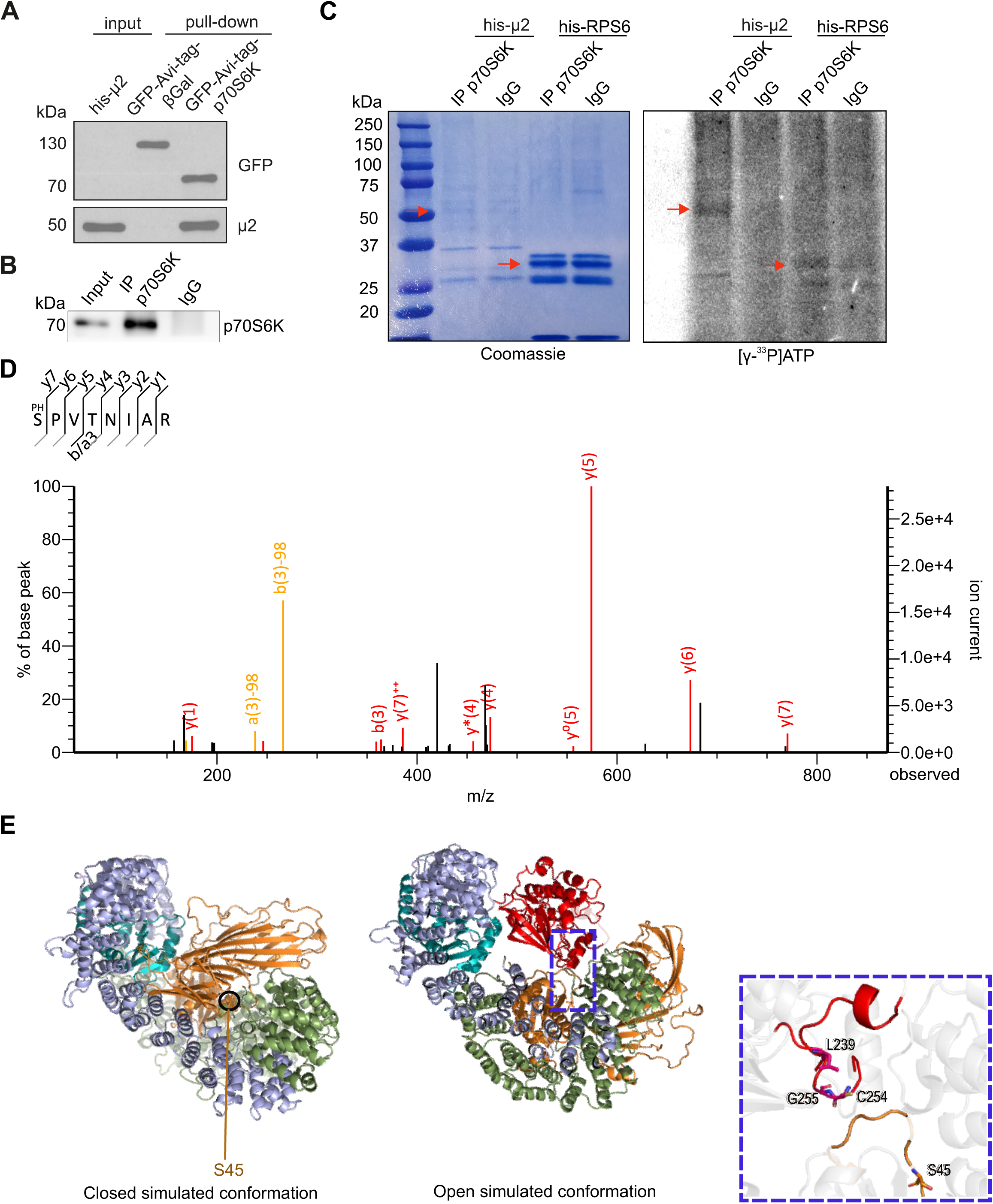
p70S6K phosphorylates μ2. **A.** Western blot evaluation of the binding assay results. Recombinant his-tagged μ2 interacts with GFP-Avi-tag-p70S6K. GFP-Avi-tagged p70S6K or β-galactosidase were expressed together with BirA in HEK293T cells, affinity-purified using M-280 streptavidin Dynabeads, incubated with recombinant his-tagged μ2 and analysed by immunoblotting. Input, 10% of lysate added to the assay. A representative example from *N=3* independent experiments. **B.** Western blot showing immunoprecipitation of p70S6K from mouse brain extract, that was used for kinase assay. Input, 10% of lysate used for immunoprecipitation. Shown is a representative example of *N= 3* independent experiments. **C.** Results of kinase assay using p70S6K immunoprecipitated from mouse brain lysates incubated with the indicated his-μ2 and C-terminal fragment of RPS6 (as positive control) in the presence of radiolabeled adenosine 5′-triphosphate (ATP). The left panel shows Coomassie R-250 staining of SDS-PAGE gels with the analyzed proteins. The right panel shows the radioactive signal level of [γ-33P] ATP incorporated into the analyzed proteins. Red arrows point to the phosphorylated recombined his-µ2 and his-RPS6. A representative example from *N* = 2 independent experiments is shown. **D.** Identification of p70S6K-dependent phosphorylation site in the μ2 subunit of the AP2 adaptor complex. Representative MS/MS fragmentation spectrum of a peptide encompassing phosphorylated serine 45 (S45) in μ2, identified after pull-down of μ2 overexpressed in HEK293T cells under control conditions (treated with DMSO). **E.** The AP2 complex as seen in our simulations in either closed or an active conformation, in a cartoon representation colored by subunits - μ2 in gold, σ2 in blue, beta in green, and sigma in cyan. Left, the AP2 in a simulated closed conformation shows S45 buried deep within the core. Right, the p70S6 kinase (red cartoon) is docked onto the AP2 core in an open conformation, indicating its accessibility for phosphorylation. The inset panel shows the active site of p70S6K and S45 in a higher magnification.

Next, we tested whether p70S6K phosphorylates μ2. Using *in vitro* ^33^P radioisotope kinase activity assay with p70S6K immunoprecipitated from mouse brain lysates (**Fig. 3B**) and purified his-tagged μ2 or the his-tagged C-terminal part of RPS6 (positive control) (**Fig. 3C**), we confirmed the ability of p70S6K to phosphorylate μ2. At the same time, our positive control (RPS6) was also phosphorylated by p70S6K, while the negative control (non-specific IgG) showed no phosphorylation of both substrates. (**Fig. 3C**). Overall, our results suggest that μ2 is a substrate for p70S6K.

To further pinpoint the residues in μ2 phosphorylated by p70S6K, we turned to a mass spectrometry-based approach. In the samples derived from control (DMSO-treated) cells, we observed the phosphorylated peptide SPVTNIAR (S45 phosphorylation, **Fig. 3D**) which was not present in the samples obtained from p70S6Ki-treated cells (**Table S1**). Of note, phosphorylation of S45 was confirmed in the PhosphoSite Plus database and has been reported in a previous study as a modification influencing the function of AP2 complex (Liu *et al*, 2019). To further confirm the possibility of μ2 phosphorylation by p70S6K in the context of the entire AP2 complex, we used the protein-protein docking approach. In the simulated closed conformation, S45 is buried deeply within the core of AP2, leaving insufficient spatial allowance for p70S6K binding (**Fig. 3E**). However, S45 becomes available for phosphorylation by p70S6K when AP2 is in its simulated open conformation (**Fig. 3E**), substantiating the notion that this phosphorylation may constitute a regulatory mechanism acting after the assembly of the functional AP2 complex. Therefore, we decided to further explore the functional links between p70S6K activity, S45-μ2 phosphorylation and the regulation of CME, and to test whether the absence of S45-μ2 phosphorylation mimics the effects of p70S6K inhibition on CME.

### Lack of S45-μ2 phosphorylation phenocopies the effects of p70S6K inhibition on transferrin internalization

To further investigate the possibility that p70S6K activity affects CME by controlling S45-μ2 phosphorylation, we tested whether the absence of this posttranslational modification has similar effects as the use of p70S6Ki described above. To this end, we performed a transferrin uptake assay in HeLa KO μ2 cells transfected with plasmids encoding myc-μ2^WT^ or myc-μ2^S45A^, a μ2 variant in which S45 has been replaced by alanine so that the protein cannot be phosphorylated at this site and mimics the properties of the unphosphorylated form. Myc-μ2^WT^ and myc-μ2^S45A^ were expressed at comparable levels and had no effect on the expression levels of endogenous β2 AP2 subunit and TrfR (**Fig. 4A, B**). Comparison of these lines showed that the uptake of fluorescently labeled transferrin was significantly decreased in μ2^S45A^ cells compared with μ2^WT^ cells (**Fig. 4C, D**). Flow cytometry-based experiments confirmed that cells expressing the μ2^S45A^ mutant are less efficient at internalizing transferrin, although this effect diminishes over time (Fig S2C-D). The difference in transferrin uptake was not due to a reduced amount of TrfR (**Fig. 4E, F**) or downregulation of TrfR surface expression reflected by the surface binding of transferrin (**Fig. 4G, H**). These results indicate that the absence of S45-μ2 phosphorylation, just like p70S6K inhibition, significantly downregulates CME.

**Figure 4.**
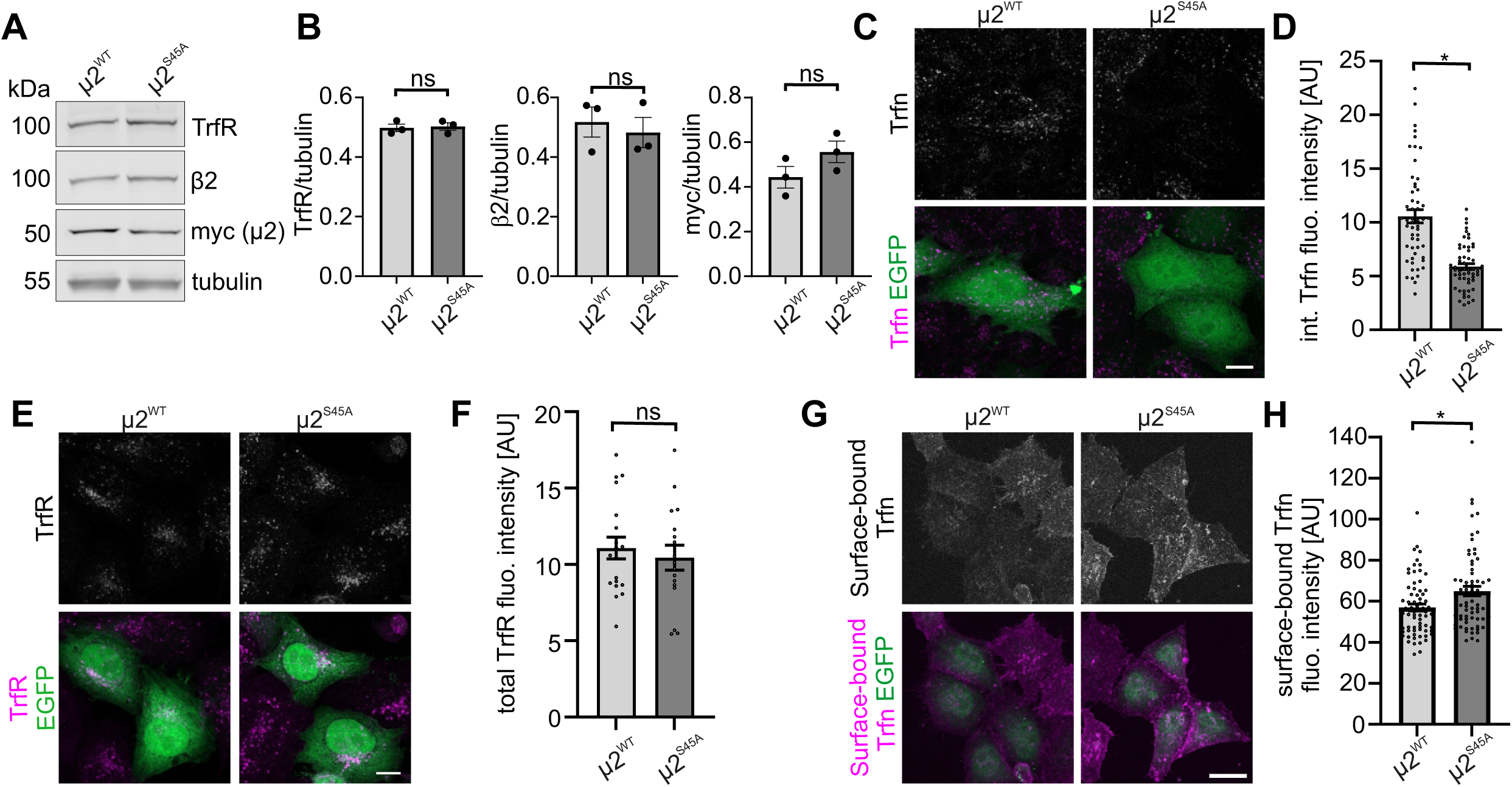
Lack of S45 phosphorylation of µ2 mimics the effects of p70S6K inhibition on transferrin internalization. **A.** Western blot analysis of TrfR, β2, myc-µ2 and tubulin in protein lysates obtained from HeLa KO µ2 transfected with a plasmid encoding myc-µ2^WT^ or myc-μ2^S45A^. **B.** Quantitative analysis of TrfR, β2, and myc-µ2 levels. *N* = 3 independent experiments. *ns* – non-significant (paired t-test; TrfR [t = 0.1878, df = 2], β2 [t = 0.3531, df = 2], myc-µ2 [t = 1.167, df = 2]). **C.** Representative images of HeLa KO µ2 transfected with plasmids encoding myc-µ2^WT^ or myc-μ2^S45A^ together with EGFP, after 5-minute uptake of Alexa Fluor 647-labeled transferrin (magenta). EGFP is shown in green. Scale bar = 10 μm. **D.** Quantification of the mean fluorescence signal intensity of internalized Alexa Fluor 647-conjugated transferrin in cells treated as in *C*. Data are presented as mean ± SEM. *N* = 3 independent experiments. Number of analyzed cells (*n*) =54 cells (µ2^WT^), 58 cells (μ2^S45A^), \*\*\**p* < 0.001 (Mann-Whitney test, U=509). **E.** Representative confocal images of HeLa KO μ2 transfected with plasmids encoding myc-μ2^WT^ or myc-µ2^S45A^ together with EGFP, immunofluorescently stained for TrfR (magenta). Scale bar = 10 μm. **F.** Quantification of TrfR fluorescence signal intensity level in cells treated as in *E*. Mean ± SEM is shown. *N* = 2 independent experiments. Number of analyzed cells (*n*) = 19 (µ2^WT^), 17 (µ2^S45A^). *ns*-non-significant, (unpaired t-test, t = 0.5813, df = 34). **G.** Representative confocal images of cell surface-bound transferrin conjugated with Alexa Fluor 647 (magenta) in HeLa KO μ2 cells transfected with plasmids encoding myc-μ2^WT^ or myc-µ2^S45A^ and EGFP (green). Scale bar = 20 μm. **H.** Quantification of the mean fluorescence intensity of transferrin conjugated with Alexa Fluor 647 bound to the cell surface in cells treated as in *G*. Mean ± SEM is shown. *N* = 4 independent experiments. Number of analyzed cells (*n*) = 68 (µ2^WT^), 67 (µ2^S45A^). \**p* < 0.05 (Mann-Whitney test, U = 1729).

### Blocking S45 phosphorylation produces mild, context-specific defects partly overlapping with apm-2 loss-of-function phenotypes

Phosphorylation of µ2 at serine 45 is required for efficient transferrin uptake in mammalian cells, but its organismal relevance has remained unclear. The serine 45 residue is conserved between human µ2 and the *C. elegans* APM-2 subunit (**Fig. S3A**). To assess the *in vivo* consequences of substituting this conserved residue, we used CRISPR-Cas9 to generate an apm-2(S45A) knock-in strain (**Fig. S3B**). Previous studies established that mutations in the AP2 α and µ subunits (*apa-2* and *apm-2*), particularly strong loss-of-function alleles, cause pleiotropic defects including reduced body length (Dpy), egg-laying defects (Egl), uncoordinated locomotion (Unc), and the distinctive ‘jowls’ phenotype with bilateral head bulges (Hollopeter *et al*, 2014; Gu *et al*, 2008, 2013).

To assess whether S45A substitution affects overall body morphology, we quantified body length and midbody width in *apm-2(S45A)* animals. S45A worms were marginally shorter and slightly wider than wild-type controls (**Fig. 5A-C**). Notably, body proportions remained within the normal range, and the characteristic ‘jowls’ phenotype was entirely absent (**Fig. 5A**). Thus, the S45A substitution is associated with subtle morphological shifts that are less pronounced than those reported for strong apm-2 loss-of-function alleles.

**Figure 5.**
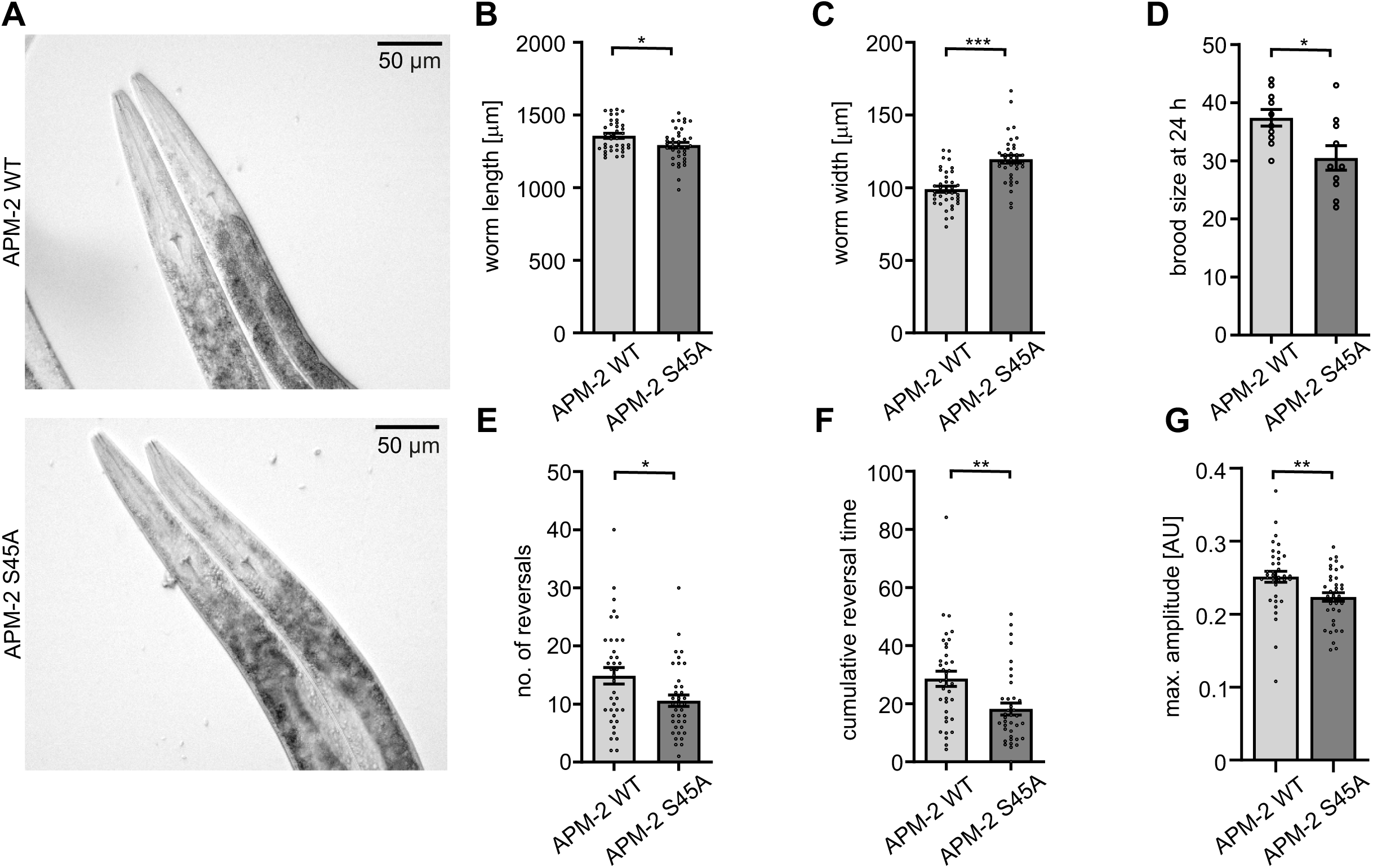
Blocking S45 phosphorylation produces mild, context-specific defects partly overlapping with *apm-2* loss-of-function phenotypes A. Representative images of young adult *C. elegans* heads: wild type (N2) and APM-2S45A mutant. Scale bar = 50 µm. **B** and **C.** Comparison of the length and width between control (WT – wild type, N2) and APM-2S45A animals. *\*p* < 0.05 (Mann–Whitney test, U = 492), *\*\*\*p* < 0.001 (unpaired t-test, t = 6.026, df = 72). **D.** The number of eggs laid by wild-type (N2) and APM-2S45A worms after a 24-hour egg-laying period. *\*p* < 0.05 (unpaired t-test, t = 2.697, df = 18). **E-G.** Number of reversals, cumulative reversal time and max. amplitude of the sinusoidal movement of wild-type (N2) and APM-2S45A worms measured by the WormLab system (MBF Bioscience). *\*p* < 0.05, *\*\*p* < 0.01 (Mann–Whitney test, [E] U = 485, [F] U = 402.5, [G] U = 399).

Given the well-documented egg-laying defects associated with strong apm-2 loss-of-function alleles, we next assessed egg-laying performance in *apm-2(S45A)* animals. Compared with wild-type controls, *apm-2(S45A)* worms laid statistically significantly fewer eggs under our assay conditions, although this effect was modest (**Fig. 5D**). In contrast, offspring survival and adult lifespan were indistinguishable from wild type (**Fig. S3C, D**), indicating that the egg-laying defect is limited in magnitude and does not reflect a general impairment of organismal viability or fitness.

Finally, we quantified several locomotory parameters in young adult *apm-2(S45A)* worms. While overall motility remained comparable to that of wild-type animals, we identified reproducible and statistically significant alterations in backward movement, including a reduced number of spontaneous reversals, shorter cumulative reversal time, and a decreased amplitude of sinusoidal body bends (**Fig. 5E-G**). These differences did not produce a gross Unc phenotype or impair forward locomotion, suggesting that the observed effects are subtle and behaviorally restricted. Taken together, our results show that in *C. elegans*, apm-2(S45A) substitution elicits a subset of AP2-associated phenotypes (e.g., in egg-laying or specific aspects of locomotion) but these manifestations are limited and substantially less severe than those associated with complete AP2 loss of function.

### Phosphorylation of S45 of μ2 influences the conformation of AP2 and is needed for the effective CME

Our results indicate that phosphorylation of S45 of μ2 contributes to efficient CME. At the same time, S45 phosphorylation does not affect the ability of μ2 to be effectively incorporated into AP2 or its interaction with TrfR. Therefore, based on available literature concerning AP2 conformational changes (Collins *et al*, 2002; Jackson *et al*, 2010; Beacham *et al*, 2018; Kelly *et al*, 2008; Kovtun *et al*, 2020; Wrobel *et al*, 2019; Beacham *et al*, 2019), we hypothesized that p70S6K-dependent phosphorylation of S45-μ2 might influence the conformation of the AP2 complex and thus the dynamics needed for efficient CME process. To verify whether phosphorylation of S45 could favor a particular AP2 conformation we performed molecular dynamics (MD) simulations of the AP2 complex (Fig. **6A**). In detail, to examine how S45 phosphorylation shapes AP2 core dynamics in the open state (PDB ID: 2XA7), we conducted 1 μs all-atom MD simulations of two μ2 phosphorylation states: unphosphorylated and P-S45. Conformational changes were monitored using the Cα–Cα distance between α-Glu142 and μ-Asn387 as a proxy for core opening (Fig. **6A**). The unphosphorylated complex served as a baseline for comparing core opening, averaging 5.49 nm over the final 500 ns. Phosphorylation at S45 resulted in the opening of the complex, with an average distance of 7.32 nm. These simulations suggest that S45 phosphorylation may promote AP2 conformational opening. In consequence, lack of S45 phosphorylation (as e.g. in S45A mutant) should disrupt the conformational changes of the AP2 complex. Indeed, MD simulations confirmed that the S45A mutant shares the properties of the unphosphorylated form and does not favor the opening of the complex (Fig. **S4A**).

**Figure 6.**
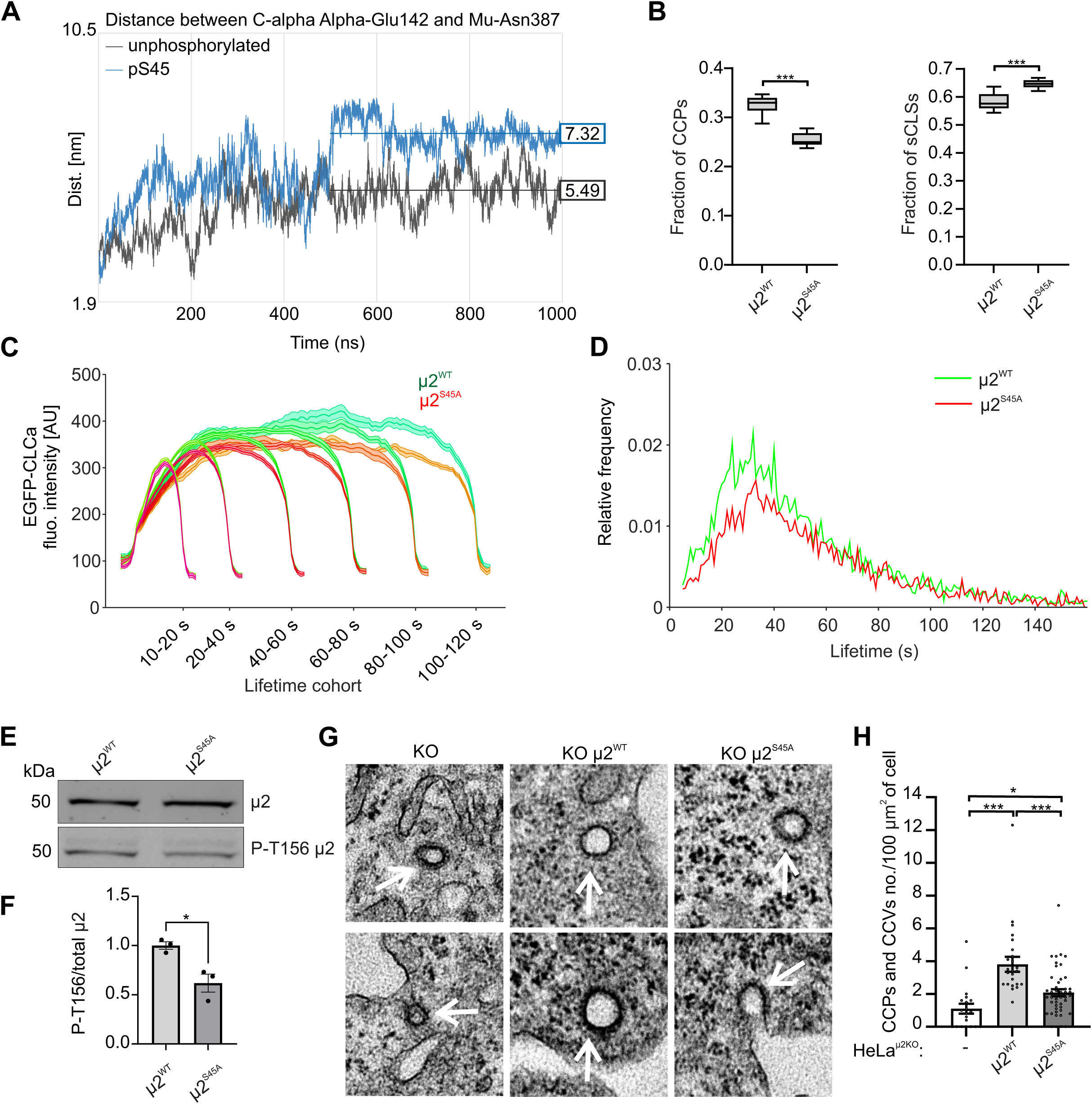
Phosphorylation of S45 of μ2 is needed for the effective formation of clathrin-coated pits. **A.** Differential effects of μ2 phosphorylation on AP2 core in open conformation. Time evolution of the distance between the C-alpha atoms of α-Glu142 and μ2-Asn387, a metric used to monitor the opening of the AP2 core, over 1000 ns all-atom MD simulations. Traces represent the unphosphorylated (black) and P-S45 (blue) states. **B.** Fractions of all detected CCPs and sCLSs for cells expressing myc-µ2^WT^ or myc-µ2^S45A^ (µ2^WT^: number of analyzed movies = 12, number of total valid tracks = 28541, µ2^S45A^: number of analyzed movies = 12, number of total valid tracks = 26194). Box plots show medians, 25th and 75th percentiles, and outermost data points. \*\*\**p* < 0.001 Matlab ranksum (*p* value 3.66e-05 for CCPs and 9.73e-05 for sCLSs). **C.** Clathrin fluorescence intensity in CCPs lifetime cohorts of cells expressing myc-μ2^WT^ (green) or myc-μ2^S45A^ (red). Intensities are shown as mean ± SE. **D.** Lifetime distributions of all CCPs found in cells expressing myc-μ2^WT^ (green) or myc-μ2^S45A^ (red). **E.** Western blot showing level of P-T156 μ2 and μ2 in HeLa KO μ2 cells stably expressing myc-µ2^WT^ or myc-µ2^S45A^. **F.** Quantitative analysis of P-T156 μ2 level. *N* = 3 independent experiments.\**p*< 0.05 (unpaired t test [t=3,851, df=4]). **G.** Electron microscopy images of CCPs in HeLa KO, HeLa KO μ2 myc-μ2^WT^ or HeLa KO μ2 myc-μ2^S45A^. CCPs and CCVs are pointed with arrows. **H.** Quantification of CCPs and CCVs number from electron microscopy images. Error bars represent ± SEM, *n* =20 cells (KO), 25 cells (KO µ2^WT^), 45 cells (KO μ2^S45A^). \*\*\**p* < 0.001; \**p* < 0.05 (Kruskal Wallis test [H=28.68] followed by multiple comparisons Dunn’s test).

If the substitution of μ2 S45 to alanine affects the conformational transitions of the AP2 complex, it should influence the dynamics of CCPs formation. To verify this assumption, we turned to the TIRF method. We constructed HeLa KO μ2 cell lines stably expressing μ2^WT^ or μ2^S45A^ together with EGFP-CLCa. Both cell lines had comparable expression levels of endogenous AP2 α and β2 subunits as well as exogenous myc-μ2 and EGFP-CLCa. Of note, after introducing exogenous μ2, expression of the endogenous μ2 was completely lost (Fig. **S4C**), which allowed us to specifically assess the effects elicited by the μ2 S45A mutant. The immunoprecipitation experiments showed that both μ2^WT^ and μ2^S45A^ were incorporated effectively into endogenous AP2 in these lines and their binding to β2 and α adaptins was similar (**Fig. S4B, C, D**). Thus, as in the case of data obtained with p70S6K inhibitor, we did not observe major differences between cells stably expressing μ2^WT^ or μ2^S45A^ in AP2 subunits expression and interaction. Considering these results, we decided to check if μ2 S45 phosphorylation might influence the efficiency and dynamics of CCP formation, which could explain the observed decrease in TrfR and PDGFR internalization seen upon p70S6Ki treatment. Thus, we compared the dynamics of CCP formation in the HeLa μ2 KO cell lines stably expressing μ2^WT^ or μ2^S45A^ together with EGFP-CLCa. In cells expressing μ2^S45A^, we observed higher fraction of sCLSs (which are not incorporated into endocytic vesicles) and lower fraction of CCPs than in the μ2^WT^ variant (**Fig. 6B**). We also observed that the matured CCPs were smaller and that the rate of clathrin polymerization was slightly reduced in 40-60 s cohort of CCPs in μ2^S45A^ cells (**Fig. 6C**), which together with an increased number of clathrin structures can be a result of disturbed CCP nucleation. At the same time cells expressing μ2^S45A^ showed decreased fraction of shorter-lived CCPs (up to 40 s) compared with the μ2^WT^ variant (Fig. **6D**). Thus, our TIRF data showing lower fraction of CCPs in μ2^S45A^ cells, together with molecular dynamics simulation results, point to the scenario where the inability to phosphorylate µ2 leads to an overall disruption of CME, potentially caused by inefficiency in obtaining or stabilizing an open conformation by AP2. If the lack of S45 phosphorylation biases AP2 toward a closed conformation, the expected effect of S45A substitution would be a reduction in the pool of μ2 phosphorylated at T156, as this post-translational modification characterizes the open state (Collins *et al*, 2002; Höning *et al*, 2005; Wrobel *et al*, 2019). In line with our prediction, the levels of μ2 phosphorylated at T156 were decreased in μ2^S45A^ cells compared to the μ2^WT^ variant (**Fig. 6E, F**).

The results obtained with TIRF pointed to less efficient CCP formation in cells in which µ2 could not be phosphorylated. To ultimately verify this observation, cells expressing μ2^WT^ and μ2^S45A^ were compared for the number and ultrastructure of CCPs and CCVs using electron microscopy. The parental line - HeLa KO µ2 - served as a negative control with diminished AP2 function. Indeed, significantly fewer CCPs and CCVs were observed in the KO µ2 cells and the cells with exogenous μ2^S45A^ expression compared to the μ2^WT^-expressing cells (**Fig. 6G, H**). In summary, we conclude that p70S6K-dependent phosphorylation of μ2 at S45 is vital for proper function of the AP2 complex and efficient formation of CCPs and, in consequence, for CME in mammalian cells (**Fig. 7**).

**Figure 7.**
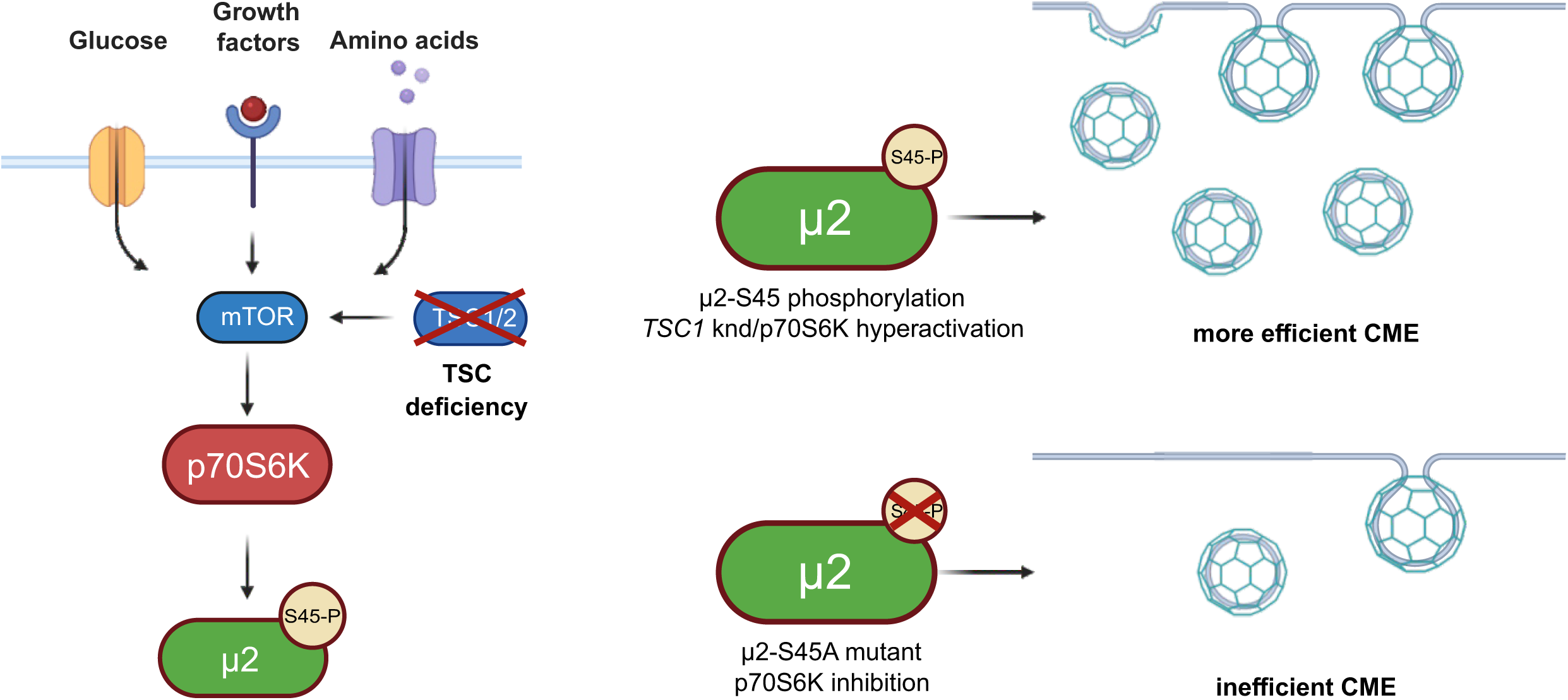
Summary of the main findings. High activity of the mTOR-p70S6 kinase pathway promotes the phosphorylation of the µ2 subunit of the AP2 adaptor complex, which is needed for efficient CME. Lack of the pathway’s activity or substitution of the wild-type µ2 with the non-phosphorylable S45A µ2 mutant alters the early steps of CME. *knd* – knockdown.

## Discussion

The role of post-translational modifications of the AP2 µ2 subunit in the regulation of CME is still not fully understood. To date, most of the studies focused on the phosphorylation at T156 (Wrobel *et al*, 2019; Partlow *et al*, 2019; Fingerhut *et al*, 2001; Ricotta *et al*, 2002), while the importance of other µ2 modifications, e.g., phosphorylation of S45, and their molecular consequences remains elusive. In this work, we have reported that p70S6K positively regulates CME by phosphorylating S45-µ2. Loss of this post-translational modification results in impaired dynamics of CCP formation and decreased CME in cells cultured *in vitro* and leads to a phenotype in *C. elegans* consistent with a partial, separation-of-function effect relative to *apm-2* loss-of-function alleles. Our results suggest that S45-µ2 phosphorylation is important for the regulation of conformational changes of the AP2 complex. Lack of this phosphorylation ultimately leads to reduced efficacy of CME, which might be caused by lack of its positive impact on promoting open AP2 conformation.

### p70S6K is µ2-S45 kinase

Several kinases have been shown to regulate CME by phosphorylating either the CME cargo or the elements of the endocytic machinery (Liberali et al, 2008). Here, we uncovered a mechanism whereby p70S6K controls CCV formation by phosphorylating µ2 subunit of the AP2 complex. The involvement of p70S6K in the control of CME is supported by a previous report showing that pharmacological inhibition or knockdown of p70S6K leads to decreased rates of albumin endocytosis and less efficient formation of CCVs in opossum kidney proximal tubule cells (Grahammer et al, 2017). This former study, however, did not explore the mechanism behind the regulatory role of p70S6K nor the relevance for other cell types, cargoes and species. Our study shows that p70S6K activity is essential for endocytosis of TrfR and PDGFR in human cells. Moreover, MS analysis pointed to the S45 of µ2 as a target for p70S6K-dependent phosphorylation. So far, the only kinase shown to phosphorylate µ2 at S45 was CDK5 immunoprecipitated from murine brain (Liu et al, 2019). Here, we show that also p70S6K immunoprecipitated from murine brain phosphorylates µ2. The possibility that the same amino-acid sequence of a protein can be phosphorylated by different kinases, whether from the same or from distinct kinase families, is not unusual. A good example is the phosphorylation of µ2 at T156. It has been demonstrated that T156 is phosphorylated not only by AP2-associated kinase 1 (AAK1) but also by BMP-2-inducible protein kinase (BMPK2) and cyclin G-associated kinase (GAK) (Conner & Schmid, 2002; Ramesh et al, 2021; Umeda et al, 2000). It remains unclear why, and under which conditions, different kinases phosphorylate µ2 at T156. For other kinases and their substrates (e.g., RPS6), such differences may depend on the expression level of a given kinase in a particular cell type, its localization or access to the phosphorylation site within specific subcellular compartments, extracellular stimuli, or metabolic state. Another important factor is whether phosphorylation requires priming i.e., prior phosphorylation of other residues by other kinases. For example, the best-characterized substrate of p70S6K1 - RPS6 - can also be phosphorylated at the same sites (S235/236) by ribosomal S6 kinase (RSK), protein kinase A (PKA), protein kinase C (PKC), protein kinase G (PKG), casein kinase 1 (CK1), or death-associated protein kinase (Biever et al, 2015; Meyuhas, 2015). Involvement of the particular kinase depends on the upstream signal: p70S6K1 is activated downstream of receptor tyrosine kinase (RTK) signaling, whereas PKA responds to G-protein-coupled receptor (GPCR) activation. Notably, phosphorylation by PKA or RSK does not require priming, i.e., prior phosphorylation of other residues (such as S240/244) by other kinases, whereas p70S6K1 does. On the other hand, the duration and dynamics of phosphorylation at the same site may also differ depending on the kinase involved. Determining whether similar principles apply to the µ2-S45 phosphorylation remains an open question and will require further studies. Finally, it should be noted that the modeling method we used does not allow us to predict the position of the AP2 complex relative to the membrane. Thus, elucidating how p70S6 kinase accesses the µ2 subunit when AP2 is attached to the plasma membrane in its open conformation requires further investigation that is beyond the scope of this study.

### The role of S45-µ2 phosphorylation in regulating CME

We found that S45-µ2 is phosphorylated in a p70S6K-dependent manner and that this modification is essential for efficient TrfR internalization. This result appears to contrast with the conclusions of Liu et al. (Liu *et al*, 2019) who showed that the presence of a short peptide competing with S45 phosphorylation enhances the internalization of EGFP-TRPV1 overexpressed in HEK cells. The reason for this difference is not entirely clear, but in our opinion, there are at least two possible explanations. First, the different cargo studied; second, the alternative approaches to inhibit S45 phosphorylation. In their work, Liu et al. analyzed the overexpressed TRPV1 receptor levels in subcellular fractions, while endogenous TrfR levels were shown as a gel loading control and appeared unchanged regardless of the manipulations applied to remove CDK5 or interfere with S45 phosphorylation. Thus, in the study by Liu et al., the internalization of EGFP-TRPV1 and endogenous TrfR appear to be differentially affected by the absence of CDK5 or diminished S45-µ2 phosphorylation, although TrfR levels and internalization were not quantified in this work, so these assumptions should be interpreted with caution. Of note, Liu et al. used an indirect approach to prevent S45-µ2 phosphorylation: the “decoy” peptide - tat S45 as a competitor for endogenous S45-µ2. Therefore, the inhibition achieved was probably acute, and the endogenous protein could still be phosphorylated, albeit to a much lesser extent. In our approach, the wild-type µ2 was replaced by the phospho-deficient variant, so the lack of µ2 phosphorylation was chronic and irreversible. Since the lack of S45 phosphorylation likely affects the preference of AP2 to remain in one of the conformations, the acute and long-term effects are presumably different. Therefore, given the significant differences in the design of the two studies, these results could be seen as complementary, jointly supporting the role of S45 phosphorylation in the regulation of CME.

### S45-µ2 phosphorylation potentially favours open conformation of the AP2

So, what could be the function of S45-µ2 phosphorylation, and how the CME defects observed in cells expressing a non-phosphorylable mutant instead of the wild-type µ2 can be explained in this context? Based on our molecular dynamics model and experimental observations, we hypothesize that the S45-µ2 phosphorylation/dephosphorylation cycle may have a function in the conformational changes of AP2. During the CME, AP2 is recruited from the cytoplasm, adopts an open (active) conformation and binds cargo, allowing clathrin coat attachment and subsequent CCV formation and cargo internalization. After CCV formation, AP2 dissociates from the CCV and transitions to a closed (inactive) conformation. Altogether, it is known that a sequential hierarchy of interactions controls AP2 activation, coordinating the early phases of CCP formation and stabilization (Kadlecova *et al*, 2017). The T156 phosphorylation/dephosphorylation cycle of µ2 is essential for the conformational changes of AP2 and cargo binding. The S45 phosphorylation/dephosphorylation cycle of µ2 may also contribute to the conformational changes of AP2. Specifically, our molecular dynamics simulations suggest that S45 can be phosphorylated in AP2 open conformation, and its phosphorylation favors this conformation, needed for efficient incorporation of clathrin into CCVs and subsequent CME. On the other hand, AP2 with non-phosphorylated S45-µ2 or containing S45A-µ2 adopts open conformation less effectively. Indeed, through quantitative analysis in living cells (TIRF microscopy), we were able to measure the distinct early stages of CME. Lower number of efficiently formed CCPs and CCVs (Fig. 6) observed in our live cell and EM imaging points to the disruption of early steps of CME in case of the µ2 S45A variant (Fig. 6), which explains the decreased internalization of TrfR or PDGFR that we observed (Fig. 2 and 4), and is in line with our modeling outcome. To sum up, our results suggest that phosphorylation of S45-µ2 is important for proper function of AP2 and for the efficient progression of CME in mammalian cells. Furthermore, our findings support a model in which S45 phosphorylation acts upstream or in parallel to the well-characterized T156 modification, contributing to AP2 conformational priming rather than directly controlling cargo binding.

### mTOR-p70S6K-AP2 axis might contribute to coupling of CME to the metabolic status of the cell

Given that p70S6K-dependent phosphorylation of S45-µ2 regulates early steps of CME, the question arises regarding the potential (patho)physiological significance of this mechanism. Since mTOR-p70S6K pathway is one of the key sensors of the metabolic status of the cell, it is tempting to speculate that the mTOR-p70S6K-AP2 axis may contribute to the coupling of CME with the current metabolic needs. Such dynamically regulated coupling is key for maintaining homeostasis and safeguarding proper cell growth, migration, and cell divisions, among others (Rahmani *et al*, 2019; Antonescu *et al*, 2014). Thus, the positive influence of the mTOR-p70S6K pathway on the initiation of CCV formation could serve to adjust the endocytosis rate to the current status of the cell.

On the other hand, the very same mechanism could also be implicated in the pathological conditions linked to mTOR and p70S6K overactivation, such as mTORopathies (Karalis & Bateup, 2021; Switon *et al*, 2017) and cancer (Wu *et al*, 2022; Zou *et al*, 2020). Contribution of dysregulated endocytosis to cancer pathogenesis is becoming increasingly recognized. Nutrient intake, signaling from cell membrane receptors, and the turnover of adhesion molecules exemplify key cellular processes that highly depend on the CME (Antonescu *et al*, 2014; Sorkin & von Zastrow, 2009; Hupalowska & Miaczynska, 2012; Miaczynska *et al*, 2004) and are at the same time crucial for oncogenesis. Thus, it has been proposed that increased metabolic demands, invasiveness, and rapid division of cancer cells are supported by dysregulated endocytic pathways (Banushi *et al*, 2023; Mellman & Yarden, 2013; Guo *et al*, 2024), and that CME can represent a target in cancer therapy (Chan & Kural, 2024). Based on our data, CME may be a potential therapeutic target not only in cancer but also in mTORopathies. The presumed involvement of dysregulated CME in supporting cell hypertrophy in these diseases is an exciting hypothesis for future studies.

## Material and Methods

### Antibodies

Commercially available primary antibodies used for this study are listed in Table 1.

**Table 1.**
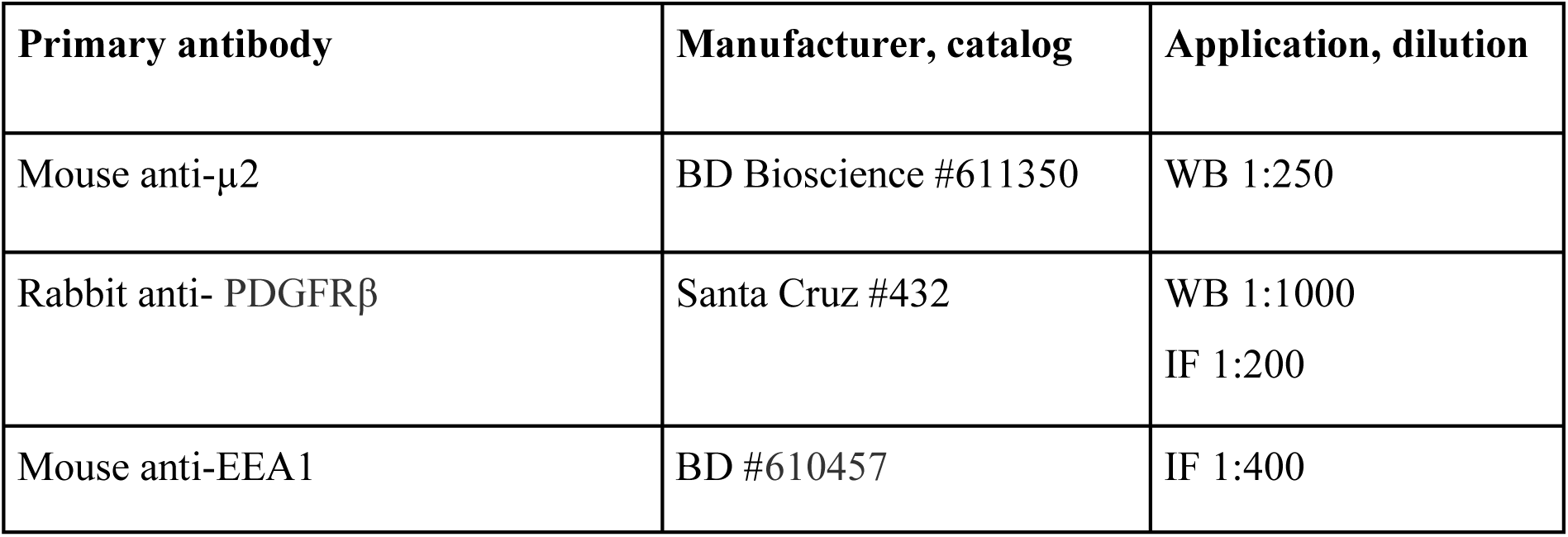
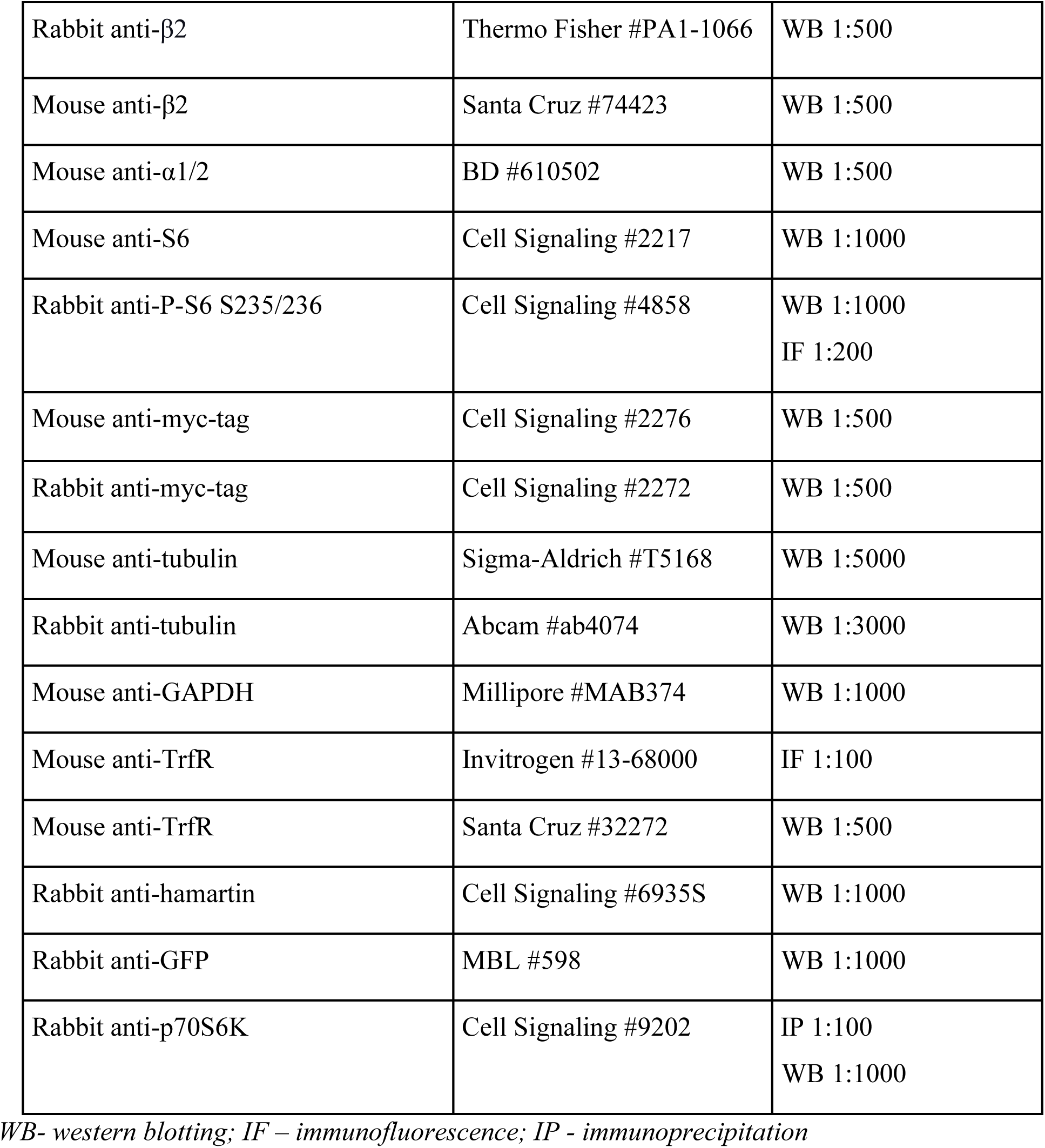
Primary antibodies used for the study.

Alexa Fluor 488-, 568-, and 647-conjugated secondary antibodies (anti-mouse, and anti-rabbit) were obtained from Thermo Fisher. Horseradish peroxidase-conjugated secondary antibodies were obtained from Jackson ImmunoResearch. Anti-mouse/anti-rabbit IRDye® 680RD and IRDye® 800CW were from LI-COR Biosciences.

### Plasmids

The following plasmids were obtained from other researchers or are commercially available: pCDH-EF1a-MCS-PGK-PURO, pVSV-G, pRSV-Rev and pCgpV plasmids (Cell Biolabs, Inc., SBI’s lentiviral system; kind gift from Krystyna Bienkowska-Szewczyk lab), pUltra (Addgene #24129), pUltra-Hot (Addgene #24130), pUltra-Chili (Addgene, # #48687), HA-BirA (de Boer *et al*, 2003) (kind gift from Casper Hoogenraad lab), pEGFPC2-Avi-tag-βGal (Swiech *et al*, 2011), pIRESneo2-myc-Ap2m1 (Motley *et al*, 2006) (kind gift from Robinson lab); p70S6K, his-RPS6 (all kind gift from Ivan Gout), pMD (Addgene: 12259; kind gift from Malcolm Moore), pPAX (Addgene: 12260; kind gift from Malcolm Moore), pGFP-CLCa (kind gift from Lucas Pelkmans), pLKO.1-TRC-shTSC1 (Pawlik *et al*, 2022). The pIRESneo2-myc-Ap2m1^S45A^ was obtained by site-directed mutagenesis of pIRESneo2-myc-Ap2m1 with use of QuikChange Site-Directed Mutagenesis System (Stratagene, Santa Clara, CA) and following primers: Ap2m1S45A_fw (5’-CGGCAGCAGGTGCGCGCCCCTGTCACAAACATCGC-3’) and Ap2m1S45A_rv (5’-GCGATGTTTGTGACAGGGGCGCGCACCTGCTGCCG-3’). pUltra-μ2^WT^, pUltra-μ2^S45A^, pUltra-Hot-μ2^WT^ and μ2^S45A^ plasmids for lentiviral vectors construction were obtained by PCR amplification of μ2^WT^ and μ2^S45A^ encoding sequences from pIRESneo2-myc-Ap2m1^WT^ and Ap2m1^S45A^ plasmids, respectively, using forward 5’ AAAATCTAGAACCGGTATGATCGGAGGCTTATT 3’ and reverse 5’ AAAAGCTAGCCTAGCAGCGGGTTTCGTAAATG 3’ primers and subsequent subcloning into XbaI and NheI sites of pUltra-Hot plasmid. pET28a-Ap2m1 was obtained by cloning rat Ap2m1 cDNA, into pET28a vector using 5’ AAGAAGCTTTAATGATCGGAGGCTTATTCATC 3’and 5’ CTCCTCGAGCTAGCAGCGGGTTTCGTAAATGC 3’.

To generate plasmid to produce lentiviral vectors encoding clathrin light chain (CLCa) in N-terminal fusion with GFP (pCDH-EF1a-MCS-PGK-PURO-GFP-CLCa), pGFP-CLCa was used as a template for PCR reaction. The reaction was performed with Q5® High-Fidelity DNA Polymerase (New England BioLabs Inc.) with following primers: forward 5’ AAAAGCGGCCGCGCCACCATGGTGAGCAAGG 3’ and reverse 5’ AAAAGTCGACTCAGTGCACCAGGGGGGCCT 3’. PCR product was subcloned into EcoRI and SalI restriction sites of pCDH-EF1a-MCS-PGK-PURO. pEGFPC2-Avi-tag-p70S6K was obtained by subcloning PCR amplified p70S6K sequences into EcoRI/SalI sites of A698 plasmid (pEGFPC2-Avi-tag). The following primers were used: forward 5’ GAAGAATTCATGGAGGCAGGAGTGTTTGACATAG 3’ and reverse 5’ GTCGTCGACTCATAGATTCATACGCAGGTGC 3’. pUltra-mCherry-shTSC1 plasmid was generated by digestion of pSuper-GFP shTSC1 vector with PstI and SalI restriction enzymes. Next, the fragment encoding the H1 promoter and TSC1 shRNA coding sequence (GAAGAAGCTGCAATATCTA) was blunted and cloned into the pUltra-mCherry vector at the SnaBI site. The TdTomato coding sequence in the vector was replaced with mCherry derived from pUltra-Hot plasmid, using SpeI/BsrGI restriction enzymes. Super-GFP shTSC1 vector was generated by annealing oligonucleotides: forward 5’ GATCCCCGAAGAAGCTGCAATATCTATTCAAGAGATAGATATTGCAGCTTCTTCTTTTTGGAAA 3’ and reverse 5’ AGCTTTTCCAAAAAGAAGAAGCTGCAATATCTATCTCTTGAATAGATATTGCAGCTTCTTCGGG 3’, followed by cloning into BglII/HindIII restriction sites of pSuper GFP.

pEGFPC2-Avi-tag-Ap2m1 was obtained by subcloning PCR amplified Ap2m1-pCDNA sequences into BglII/HIndIII sites of A698 plasmid (pEGFPC2-Avi-tag). The following primers were used: forward 5’ AGAAGATCTCTATGATCGGAGGCTTATTCATC 3’ and reserved 5’ AAGAAGCTTCTAGCAGCGGGTTTCGTAAATGCC 3’. The plasmids generated in this study are available from the corresponding authors upon reasonable request.

### Generation of lentiviral vectors and cell lines

Lentiviruses were generated by transfection of HEK293T cells with calcium phosphate method (CalPhos™ Mammalian Transfection Kit, Takara) for 48 hours. After this time supernatants were collected and used for transduction of HeLa cells wild type or HeLa KO μ2. First, we generated and sorted cell lines expressing EGFP-CLCa, following transduction with lentiviruses carrying μ2 constructs or TSC1 shRNA. All generated cell lines were sorted for the same level of EGFP and mCherry fluorescence between variants using BD FACSAria II cell sorter (Becton Dickinson) using blue laser (488 nm, 13 mW) and 530/30 and 616/23 nm emission filters.

### C. elegans strains

Worms were maintained on nematode growth medium (NGM) plates seeded with OP50 *Escherichia coli* bacteria at 20°C unless otherwise stated (Brenner, 1974). The amp-2 gene mutant (c. 375 T>C. p.S45A) was generated using CRISPR/Cas9 method as previously described (Dokshin *et al*, 2018). The crRNA sequence used was TCGCTCGCCAGTCACCAACA. The *apm-2* gene locus was sequenced, and mutation confirmed. The resulting *amp-2(WOP370)* was backcrossed to wild-type N2 worms.

### Drugs treatment

p70S6K inhibitor (Calbiochem, PF-4708671) was used in 10 µM concentration. p70S6Ki was dissolved in DMSO (Sigma-Aldrich, D8418), whose final concentration in the culture medium did not exceed 0.1%.

### Cell line cultures and transfection

HeLa cells and HeLa KO µ2 cells were a kind gift from Juan Bonifacino laboratory. HEK293T cells and CCD-1070Sk (CRL-2091) human normal foreskin fibroblasts were purchased from ATCC. HEK293T and HeLa cells were grown in Dulbecco’s modified Eagle’s medium (DMEM) that contained 10% fetal bovine serum and 1% penicillin-streptomycin (all from Sigma-Aldrich). CCD-1070Sk cells were cultured in Minimum Essential Medium (MEM) supplemented with 10% fetal bovine serum (FBS), and 2 mM L-glutamine (all from Sigma-Aldrich). All cells were maintained at 37°C in a 5% CO_2_ atmosphere and PCR-based tests for mycoplasma contamination were routinely performed. For experiments with use of confocal or TIRF microcopy, cells were grown on precision cover glasses (Marienfeld, 0107032). HEK293T and HeLa cells at 70% confluency were transfected using polyethylenimine PEI 25K™ (Polysciences, 23966) according to the manufacturer’s protocols. After transfection cells were grown in DMEM supplemented with 5% FBS for 48 h. For flow cytometry experiments, HeLa KO µ2 cells were transfected using the same protocol. However, in these experiments, cells were cultured on plastic in 24-well plates instead of on glass coverslips.

### Immunofluorescence

HeLa KO µ2 cells were fixed with 4% PFA (containing 4% sucrose in PBS pH 7.4) for 10 min and washed three times with PBS. After fixation cells were blocked for 1 hour in PBS containing 5% donkey serum and 0.3% Triton X-100. Primary antibodies were incubated in PBS containing 1% BSA and 0.3% Triton X-100 at 4°C overnight. After overnight incubation coverslips were washed three times with PBS. Alexa 488-and Alexa 594–conjugated secondary antibodies were used for 1 hour at room temperature. Coverslips were washed three times with PBS and mounted with Prolong Gold with DAPI (Thermo Fisher, P36941).

### Transferrin uptake and PDGFR internalization assay

Cells were starved at 37°C in a 5% CO_2_ atmosphere in non-supplemented DMEM medium for 18 h. Alexa Fluor 647-conjugated transferrin from human serum (Thermo Fisher, T23366) was then added to the cells for 5 min at a final concentration of 25 μg/ml. The cells were then rinsed with medium, fixed with 4% paraformaldehyde (containing sucrose), and washed 3 times with PBS. PDGFR internalization by CCD fibroblasts and results quantification were performed as described in Jastrzebski et al. (Jastrzębski *et al*, 2017). In brief, cells were treated with PDGF (PeproTech #100-14B; 50 ng/ml) for 15 minutes and fixed with 4% paraformaldehyde prior to staining with anti-EEA1 and anti-PDGFR antibodies and image acquisition.

For transferrin surface staining HeLa KO μ2 cells after starvation for 18 h were incubated for 15 min on ice following 15 min incubation on ice with transferrin conjugated with Alexa Fluor 647 (25 μg/ml) in DMEM without FBS. Next, cells were washed with cold PBS and fixed with cold 4% paraformaldehyde for 15 min on ice and 15 min at room temperature, followed by washing 3 times in PBS.

### Fixed cells image acquisition and analysis

Images of fluorescently labeled *in vitro* cultured cells were acquired with a Zeiss LSM800 confocal microscope (40· or 63· objective) with the settings kept constant for all scans in a series. Z-stacks (interval 0.2 µm – 0.5 µm) were converted to single images using a maximum intensity projection. The mean fluorescence intensity was measured using *ImageJ,* except for PDGFR internalization analysis that was performed as described previously (Jastrzębski *et al*, 2017). In brief, the intensity of PDGFR staining within EEA1-positive structures was quantified. For each experiment and condition, 10-20 fields of view were analyzed.

### Transferrin uptake and flow cytometry measurements

Cells were either treated with 10 µM p70S6K inhibitor for 2 h or with 0.1% DMSO as a vehicle control (HeLa cells), or HeLa μ2 KO cells were transfected with pULTRA-μ2^WT^ or pULTRA-μ2^S45A^ plasmids 48 h prior to the experiment, as described above. Cells were starved at 37°C in a 5% CO2 atmosphere in non-supplemented DMEM medium for 18 h. Prior to the uptake assay, plates with cells were placed on ice for 5 min. Cells were then incubated with serum-free medium containing 15 µg/mL Alexa Fluor 647-conjugated transferrin from human serum (Thermo Fisher, T23366) for 5 min on ice. For the 0 min time point, cells were maintained on ice throughout the experiment and were not transferred to 37°C. For internalization assays, cells were shifted to 37°C for 5, 10, or 15 min to allow transferrin uptake. At the end of the incubation period, plates were immediately transferred onto ice to stop endocytosis. Cells were washed once with ice-cold PBS and incubated for 1 min on ice with an acid wash solution consisting of 0.1 M sodium acetate and 0.2 M NaCl (pH 5.5) to remove surface-bound transferrin. Subsequently, cells were washed with ice-cold PBS followed by a wash with PBS at room temperature. Cells were detached by incubation with pre-warmed trypsin for 2 min. Trypsinization was stopped by adding ice-cold DMEM, and cells were gently resuspended. Cell suspensions were transferred to flow cytometry tubes and maintained on ice until analysis. Then cells were incubated for 5 min with viability dye (Via Krome 405 Fixable Viability Dye, no C36614 from Beckam Coulter). Then samples were immediately measured using CytoFLEX LX cytometer equipped with 405 nm (80 mW) violet laser and 450/45 bandpass emission filter for viability dye, 488 nm blue laser (50 mW) and 525/40 emission filter for GFP, 638 nm (50 mW) red laser and 660/20 emission filter for TrnfAF674. At least 5000 single and live cells were recorded for each condition. In case of cells transfected with pULTRA-μ2^WT^ and pULTRA-μ2^S45A^ plasmids, gating for GFP+ cells was implemented. Gating strategy is shown on Fig. S2. Transferrin uptake was quantified as the median fluorescence intensity (Median R660-APC-A). For pULTRA-μ2^WT^-and pULTRA-μ2^S45A^-transfected cells, the Median R660-APC-A was calculated for the live GFP-positive population, whereas for inhibitor-treated HeLa cells the Median R660-APC-A was calculated for the live cell population. For each experimental set, values obtained at 5, 10, and 15 min were normalized to the corresponding 0 min control.

### TIRF microscopy

Cells (3×10^5^) were seeded on coverslips coated with matrigel 2-3 hours before imaging. The cells were imaged on the Andor Revolutions XD imaging platform equipped with Zeiss TIRF illuminator, Chroma dual band 488/561 TIRF filter set, Zeiss 100x/1.46 Plan-Apo objective, Optovar magnification changer set at 1.6x and LucaR EM-CCD camera (Andor). Time-lapses were acquired using IQ3 software (Andor) at 1 Hz for 5 minutes. Detection and lifetime analysis of clathrin structures was performed with the publicly available cmeAnalysis software [25] according to the software manual. Only diffraction-limited, spatially confined GFP-containing structures that created valid tracks after tracking were included in the analysis. Valid tracks were further classified as sCLSs and CCPs based on dynamic parameters and intensity threshold in the cmeAnalysis software. Descriptive statistics of populations (1 n = 1 movie that typically contained 1-3 cells in the field of view) were shown as box plots (boxes indicate the quartiles and whiskers extend to extreme data points). Significance of differences between populations was tested with the ranksum test (Matlab).

### Electron Microscopy sample preparation, processing and imaging

Electron microscopy was performed as described earlier (Wróbel *et al*, 2022). Briefly, cells were grown on coverslips (Thermanox plastic coverslips; NUNC Brand Product) on 24-well plate. 3 h after seeding, cells were fixed in 2.5% glutaraldehyde for 2 hours, followed by 3 washings with PBS. The material was then postfixed with 1% osmium tetroxide for 1 h, washed with water, and stained in 1% aqueous uranyl acetate overnight at 4°C. In the next step, cells were dehydrated with increasing concentrations of ethanol and embedded with epoxy resin (Sigma Aldrich, #45-359-1EA-F). Resin-infiltrated samples were then incubated at 60°C for 72 h for polymerization. Polymerized blocks with cells were trimmed and cut with a Leica ultramicrotome (EM UC7) for ultrathin sections (65 nm thick). Sections were placed on nickel grids, mesh 200 (Agar Scientific, AGG2200N). The material was examined with a transmission electron microscope Tecnai T12 BioTwin (FEI, Hillsboro, OR, USA) equipped with a 16-megapixel TemCam-F416 (R) camera (TVIPS GmbH).

### Electron Microscopy images quantification

Images of the cells captured at 13.000 x magnification were assembled to have continuous cytosol ultrastructure. CCPs and CCVs, visualized with electron microscopy, were manually recognized and counted. The area of the cell was analyzed in *ImageJ*, from micrographs of lower magnification (2.900), allowing to fit the whole cell on one or two pictures. Obtained values were then collected and analyzed in Excel software.

### Whole-cell lysate preparation

For Western blot analysis of proteins in whole cell lysate, cells were lysed in the lysis buffer (20 mM Tris pH 7,5; 150 mM NaCl; 2 mM EDTA; 0.5% Triton X-100; 0.5% NP-40; 2 mM MgCl_2_; 10% glycerol; protease and phosphatase inhibitors). For total protein concentration in whole-cell lysates, Pierce BCA Protein Assay Kit (Thermo Fisher, 23225) was used according to the manufacturer’s instructions. Then 4x Laemmli sample buffer was added to lysates followed by boiling for 5 min at 95°C.

### Production and purification of recombinant proteins in bacteria

The genes encoding C-terminal fragment of RP6S and µ2 were cloned into the pET28a vector to enable the expression of proteins carrying a C-terminal 6xHis tag. The resulting constructs were then transformed into *E. coli* BL21 or Rosetta (Novagen), respectively. The strains obtained were used for production of his-µ2 and his-RPS6 proteins. Overnight cultures (37°C, shaking) were diluted 1:100 in fresh medium (Luria Broth) and grown until an OD600 of 0.4. Protein expression was induced with 0.25 mM isopropyl β-d-1-thiogalactopyranoside (IPTG; Sigma, I6758). His-RPS6 was produced for 3 h at 37°C, whereas the his-µ2 culture was incubated overnight at 18°C after induction. The cultures were then harvested by centrifugation at 8,000 x *g* for 8 min. Cell pellet containing his-RPS6 was resuspended in a lysis buffer composed of 300 mM KCl, 50 mM KH_2_PO_4_ and 10 mM imidazole, whereas the pellet containing his-µ2 was resuspended in 50 mM NaH_2_PO_4_/Na_2_HPO_4_ (pH 8.0), 300 mM NaCl and 5 mM imidazole. Both buffers were supplemented with lysozyme (1 mg/ml) and a protease inhibitor cocktail (cOmplete EDTA-free; Roche, 11836170001). After resuspension and a 20 min incubation at room temperature, cells were lysed on ice by sonication using a Sonics VCX130 PB sonicator (Vibra-Cell). The lysates were centrifuged at 13,000 x *g* for 15 min, and the resulting supernatants were filtered through a 0.22 μm membrane to remove any debris.

The supernatant containing his-RPS6 was incubated with ProfinityTM IMAC Ni-Charged Resin (BioRad, 1560133) in a ratio of 0.2 ml of resin per 1 ml of supernatant for 1 h at 4°C with gentle end-to-end rotation. The resin was then transferred to a column, washed with lysis buffer, and the bound protein was eluted with a buffer containing 300 mM KCl, 50 mM KH_2_PO_4_ and 250 mM imidazole. The supernatant containing his-µ2 was loaded onto EconoFit Nuvia IMAC columns (BioRad, 12009285), and the protein was eluted with an imidazole gradient on the NGC chromatography system (BioRad). The eluted fractions were desalted with a use of Econo-Pac 10DG Desalting Columns (BioRad, 7322010) into storage buffer (150 mM NaCl, 50 mM Tris pH 7.5, 10% glycerol, 10 mM β-mercaptoethanol) and concentrated as needed using Amicon® Ultra Centrifugal Filters (10 kDa MWCO; Millipore, UFC8010).

### Kinase assays

Mouse brain extracts in lysis buffer (20 mM Tris pH 7.5, 150 mM NaCl, 1 mM CaCl_2_, 1 mM MgCl_2_ supplemented with protease and phosphatase inhibitors) were immunoprecipitated with an anti-p70S6K antibody at 4°C overnight. The immunoprecipitates were washed four times with washing buffer (20 mM Tris pH 7.5, 150 mM NaCl, 1 mM CaCl_2_, 1 mM MgCl_2_ supplemented with 0.1% CHAPS) and twice with kinase buffer [20 mM Tris (pH 7.5), 20 mM MgCl_2_, 1 mM EDTA, 1 mM EGTA, and 0.1 mM dithiothreitol]. *In vitro* phosphorylation assay was performed for 20 min at 30°C in a kinase buffer in presence of 2.5 μCi [γ-^33^P]ATP (Hartmann Analytic GmbH, Braunschweig, Germany) and 2.5 mM “cold” ATP using recombinant protein his-tagged µ2, or his-tagged C-terminal fragment of RPS6 (as a positive control) as substrates. The reaction was stopped by adding 4x Laemmli sample buffer and boiling for 5 min at 95°C. Next, samples were separated by SDS-PAGE. The gel was stained with Coomassie Brilliant Blue and dried. The radioactive signal was detected and analyzed using a Typhoon Trio+ Phosphorimager (GE Healthcare).

### Cell surface biotinylation assays

The assay was performed according to the manufacturer’s protocol Pierce Cell Surface Biotinylation and Isolation Kit (Thermo Fisher, A44390).

### Avi-tag pull down of in vivo biotinylated proteins and his-µ2 binding

HEK293T cells co-expressing BirA together with Avi-tag proteins were harvested and lysed on ice in lysis buffer for 15 min. M-280 streptavidin Dynabeads (Thermo Fisher, 11205D) were washed with lysis buffer (20 mM HEPES, pH 7.5, 120 mM KCl, 0.3% CHAPS supplemented with protease and phosphatase inhibitors). The cell lysate was added to the previously prepared M-280 streptavidin Dynabeads and incubated together for 1 h at 4°C while rotating. Then, beads were washed on ice four times in washing buffer (20 mM HEPES, pH 7.5, 500 mM KCl, 0.1% CHAPS) and incubated with 0.2 mg/ml his-µ2 in binding buffer: 20 mM HEPES, pH 7.5, 300 mM KCl, 0.2 µg/µl BSA and 0.1% CHAPS) for 2 h at 4 °C rotating. Finally, beads were washed four times with binding buffer, eluted in Laemmli sample buffer and analysed via SDS–PAGE and subsequent immunoblotting.

### MS/MS Analysis of Protein Phosphorylation

To identify phosphorylated residues within the µ2 sequence, GFP-Avi-tag-µ2 protein was overexpressed in HEK293T cells. Next cells were treated overnight with DMSO or p70S6Ki. GFP-Avi-tag-µ2 was purified as described above, denatured, and subjected to SDS-PAGE. The gel was stained with silver, and bands that corresponded to µ2 were excised. The samples were then subjected to “in-gel digestion” with trypsin and obtained peptides were analyzed by mass spectrometry as described previously (Gozdz *et al*, 2017) with minor modifications. Database searching was performed using the Mascot search engine (Matrix Science, London, UK; Mascot Server version 2.8.3) against the UniProtKB/Swiss-Prot *Rattus norvegicus* database (8,234 sequences) and the cRAP database (115 sequences; 38188 residues). To reduce mass errors, the peptide and fragment mass tolerance settings were established separately for individual LC-MS/MS runs (Malinowska *et al*, 2012). Enzyme specificity was set to trypsin. The mass calibration and data filtering were carried out with MScan software (http://proteom.ibb.waw.pl/mscan/). The Decoy Mascot functionality was used for keeping FDR for peptide identifications below 1%. All peptides with q-values > 0.01 were removed from further analysis. The list of identified peptides for AP2 protein (P84092) was exported to Excel.

### Western blotting

Protein samples were analyzed by SDS–PAGE electrophoresis and immunoblotting according to standard laboratory protocols. Primary antibodies used for Western blotting are listed in Table 1. For protein separation, 10% polyacrylamide gels were run at 90 V for about 20 min and then 120 V until the loading buffer reached the bottom of the gel. Proteins were transferred to a nitrocellulose membrane (Amersham™ Protran, Sigma GE10600002) using a wet blot transfer system (Bio-Rad Laboratories) for 90 min at 100 mA. Membranes were blocked for 1 h with 5% non-fat milk in TBS-T (tris-buffered saline with 0.05% Tween 20). Primary antibodies were incubated overnight at 4°C. After washing three times with TBS-T, the secondary antibody (1:10000 in 5% TBST-milk, Jackson ImmunoResearch) was incubated for 1 hour at room temperature. The signals were detected using ECL solution or ImageStudioLite (LI-COR). In case of fluorescent signal detection in LI-COR system, TBS was used in all steps instead of TBS-T.

### Immunoprecipitation

For the immunoprecipitation (IP) of µ2 with myc tag from HeLa KO µ2, cells stably expressing EGFP-CLCa and myc-µ2^WT^ or myc-µ2^S45A^ were seeded on 10 cm diameter dish (5×10^6^ cells/dish). After 24 h cells were washed 2 times with cold TBS buffer, harvested and lysed on ice for 15 min in lysis buffer (20 mM Tris pH 7.5, 150 mM NaCl, 0.5% CHAPS (3-((3-cholamidopropyl) dimethylammonio)-1-propanesulfonate) supplemented with protease and phosphatase inhibitors. The lysates were incubated with Pierce™ Anti-c-Myc Magnetic Beads (ThermoFisher Scientific, #88843) with rotation for 2 hours at room temperature. After this time beads were washed 4 times with washing buffer (20 mM Tris pH 7.5, 300 mM NaCl, 0.1% CHAPS), eluted in 2x Laemmli sample buffer, and analyzed by Western blot.

### Worm mobility and size

Approximately 10 young adult worms per replicate were placed onto NGM plates and were recorded for 2 min using the WormLab system (MBF Bioscience). The frame rate, exposure time, and gain were set to 7.5 frames per second, 0.0031 s, and 1, respectively. The track length and size of the individual worm were analyzed using the WormLab software (MBF Bioscience). The assay consisted of three independent biological replicates. 37-39 worms were recorded for one biological replicate.

### Lifespan assay

Lifespan measurements were done from the young adult stage at 20° C on NGM plates containing 400 µM FUdR. During lifespan measurements, worms were scored daily for movement and pharyngeal pumping until death. Animals that crawled off the plate or exhibited baggy phenotype were censored from the experiment. The essay consisted of three independent biological replicates, with each replicate comprising 20 worms.

### Egg-laying and hatching analysis

Age-synchronized hermaphrodites at the L4 developmental stage were picked for the experiment. Worms were cultured as single worms per 60 mm NGM plate, seeded with *E. coli* OP50. The assay consisted of three independent biological replicates, with each replicate comprising 10 worms. The experiment involved a 24-hour observation period during which we recorded the total egg production. Concurrently, we monitored the emergence of larvae to determine the success rate of hatching.

### In silico analyses

To investigate the effect of phosphorylation on the conformational state of the AP2 core, all-atom molecular dynamics (MD) simulations were performed using the experimental crystal structures of the AP2 core in its open (PDB ID: 2XA7) and closed (PDB ID: 4UQI) conformations.

Post-translational modification and amino acid substitution were introduced by first generating models of the full-length μ2 chain containing P-S45 modification or S45A substitution using the AlphaFold3 webserver (server was accessed on 7th July 2025) (Abramson *et al*, 2024). The coordinates of the modified residue and its local environment were then structurally aligned and superimposed onto the experimental crystal structure. The unmodified crystal structure was used for the unphosphorylated control simulations. Missing loops were modeled using MODELLER 10.6 (Eswar *et al*, 2006) via the UCSF ChimeraX interface. For the open state, missing loops comprised residues 136-141 and 222-235 in the μ2 subunit, whereas for the closed state they comprised residues 583-617 in the β subunit and residues 142–158, 223–239, and 256–260 in the μ2 subunit.

The three open-state systems (unphosphorylated, P-S45; S45A) and the closed-state unphosphorylated system weresolvated in a truncated octahedral box of OPC water with a 10 Å buffer and neutralized with counterions. The closed-state system underwent an initial 10,000-step vacuum minimization prior to solvation to relieve steric clashes introduced during loop modeling

All simulations were performed using the AMBER 24 suite with the sander module. The systems were described using the ff19SB force field for the protein, phosaa10 parameters for phosphorylated residues, the OPC water model, and the ionslm_126_opc ion parameters. Bonds involving hydrogen atoms were constrained using the SHAKE algorithm.

The systems were subjected to a two-stage minimization: first, the solvent and ions were minimized for 10,000 steps with positional restraints of 20.0 kcal/mol/Å² on all solute atoms, followed by a 10,000-step unrestrained minimization of the entire system. Following minimization, each system was gradually heated from 100K to 300K over 500ps in the NVT ensemble with weak restraints on the solute. The systems were then equilibrated for 500ps in the NPT ensemble at 300K and 1atm using a Monte Carlo barostat to achieve the correct density. A series of three 200ps equilibration steps in the NVT ensemble were then performed, with the force constant on the protein backbone gradually reduced from 10.0 to 5.0 to 1.0 kcal/mol/Å². After an additional 2 ns of unrestrained equilibration in the NVT ensemble, a 1-µs production simulation was carried out for each system in the NPT ensemble at 298K and 1atm. Throughout the protocol, temperature was maintained using a Langevin thermostat.

To quantify the conformational changes of the AP2 core, the Cα distance between α-Glu142 and μ-Ser387 was calculated across each 1-µs trajectory using the cpptraj module of AmberTools. The average distances were calculated over the final 500 ns (i.e. once the systems were deemed stabilized) to analyze the effect of phosphorylation on the openness of the core.

To model the interaction between p70S6K and the AP2 core, protein-protein docking was performed using the HADDOCK 2.4 webserver (server was accessed on 15th September 2025) (Honorato *et al*, 2024, 2021). The starting structure for human p70S6K was obtained from the Protein Data Bank (PDB ID: 3A62). The structure for the AP2 core used for docking was derived from a representative snapshot at 800 ns of the open-state molecular dynamics’ trajectory, which corresponded to the most structurally stable and open state observed. Ser45 of the μ2 chain was specified as an active (i.e. target) residue for the substrate AP2. For the p70S6K enzyme, the active (i.e. target) residues were defined as Cys254, Gly255, and Leu239 (the DFG+1 position) (Chen *et al*, 2014). These solvent-exposed residues are located on a loop adjacent to the canonical catalytic motif (Tyr216, Arg217, Asp218) (Gizzio *et al*, 2024), which was partially occluded in the crystal structure; this choice was based on the hypothesis that the active site region would become more accessible upon substrate binding. For both proteins, passive residues were automatically defined by the HADDOCK server as all residues within a 6.5 Å radius of the active residues. From the 200 water-refined models generated, the best 40 structures were grouped into 8 clusters. The final representative model was selected for analysis based on a consensus of favorable HADDOCK scores, Z-scores, cluster size, and visual inspection of the top-ranking models in PyMOL (The PyMOL Molecular Graphics System, Version 2.4, Schrödinger, LLC.). For structural comparison, a representative snapshot at 600 ns of the closed state trajectory was used to illustrate the buried position of μ2-Ser45.

## Statistical analysis

The exact number of repetitions of each experiment (N) and the number of cells (n) that were examined for the respective experiments are listed in the figure legends. The statistical analysis was performed using GraphPad Prism 10 studio or Matlab. The exact statistical test that was used in the particular analysis is provided in the figure legend. All reported *p* values were derived from two-tailed statistical tests.

## Acknowledgments

The authors thank all researchers listed in Materials and Methods for sharing reagents. We are also grateful to Prof. Juan Bonifacio for hosting AT in his lab and advising her on research directions, Volker Haucke, Antoni Wrobel, Martin Lehmann, Jan Brezovsky, Marcel Mettlen, Zhiming Chen, and Carlos Eduardo Sequeiros Borja for helpful discussions. We also thank Pankaj Thapa, Malgorzata Urbanska, and Roberto Pagano for help with experiments, and Alina Zielinska for technical assistance and support, Marcelina Firkowska, Marek Sarnacki and Angelika Jocek for laboratory management logistics. We would like to thank the Institute of Genetics and Biotechnology, Faculty of Biology, University of Warsaw, for providing access to the isotope laboratory and for technical support in the kinase assays.

This research was supported by Polish National Science Centre Preludium grant no. 2017/25/N/NZ3/01280 to AT. AT was partly financed by Polish National Science Centre Opus grants no. 2016/21/B/NZ3/03639 and 2017/27/B/NZ3/01358 to JJ. Work of AB was partly financed from Polish National Science Centre Opus grants no. 2016/21/B/NZ3/03639. JJ was partly financed by the TEAM grant from the Foundation for Polish Science (POIR.04.04.00-00-5CBE/17-00). AF was financed by Polish National Science Centre Opus grant no. 2020/39/B/NZ2/01301. The work of EL was financed from IIMCB statutory funds. The work of KM was partly financed from Polish National Science Centre Maestro grant 2020/38/A/NZ3/00447 to JJ. ARM was partly financed by I.3.4 Action of the Excellence Initiative - Research University Programme at the University of Warsaw. This research was also possible thanks to the IIMCB IN-MOL-CELL Infrastructure funded by the European Union – NextGenerationEU under National Recovery and Resilience Plan. IN-MOL-CELL Infrastructure was also funded by the European Union under Horizon Europe (Project 101059801 - RACE) and by RACE-PRIME project carried out within the IRAP programme of the Foundation for Polish Science co-financed by the European Union under the European Funds for Smart Economy 2021-2027 (FENG). We also acknowledge Polish high-computing infrastructure PLGrid (HPC Center: ACK Cyfronet AGH) for providing computer facilities and support within computational grant no. PLG/2025/017952.

## Data availability

All data generated or analyzed during this study are included in this published article and its supplementary materials. Raw data from all quantitatively analyzed experiments, as well as plasmids generated in this study, are available from the corresponding authors upon reasonable request. All software used in this study is publicly available. No custom software was developed for this work. The raw mass spectrometry data and processed results have been deposited in the jPOST repository under accession number JPST004745.

## Author contributions

AT, AB, TW, MWG, WP, JJ and ARM designed the experiments. AT, AB, TW, AF, KO, KJ, EL, KM1, KM2, MM1, ASz, ES, AM, AG, AW, AŁ, MHM, KO, JJ and ARM performed the experiments. AT, AB, TW, KO, KJ, EL, MM1, ES, AM, MM2, MWG, WP, JJ and ARM analyzed the data. AT, AB, WP, JJ and ARM wrote the manuscript. All authors read and approved the manuscript.

## Declaration of interests

The authors declare that they have no conflict of interest.

## SUPPLEMENTARY INFORMATION

**Figure S1.** mTORC1 signaling and PDGFR localization in TSC1-deficient cells.

**Figure S2.** Evaluation of transferrin uptake by flow cytometry.

**Figure S3.** Serine 45 in APM-2 in *C. elegans*.

**Figure S4.** The impact of the µ2 S45A mutation on the AP2 complex and the level of CLCa.

**Figure S5.** Blot transparency.

**Table S1.** Comparison of detected μ2 peptides and post-translational modifications between DMSO and p70S6Ki-treated samples.

---

**Figure S1.**
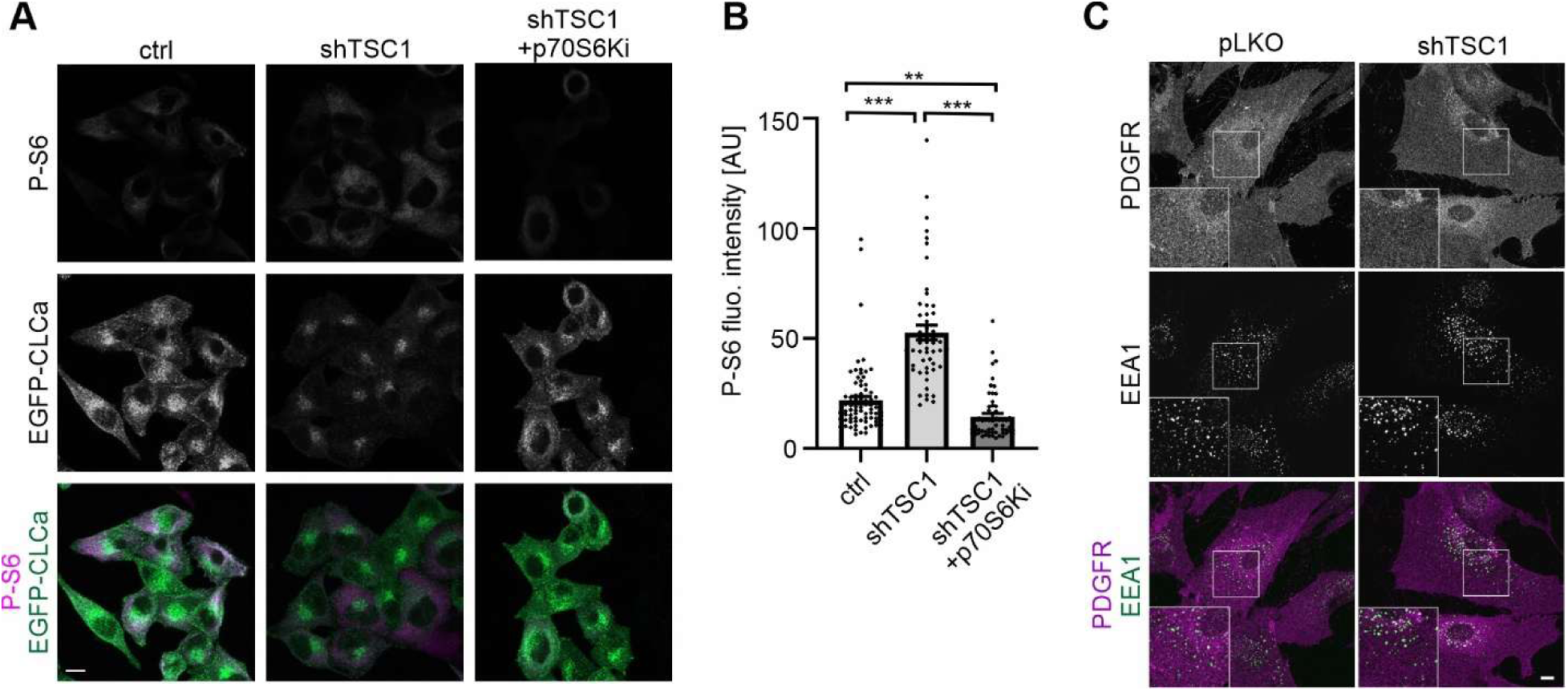
mTORC1 signaling and PDGFR localization in TSC1-deficient cells.A. Representative confocal images of immunofluorescent labeling of P-S6 (S235/236) (magenta) in HeLa cells stably expressing EGFP-CLCa (green) alone or together with TSC1 shRNA. Cells were serum-starved and treated for 2 h with 0,1% DMSO or 10 µM p70S6Ki. Scale bar = 10 µm **B.** Quantification of immunofluorescence signal of P-S6 (S235/236) normalized to cell area in cells treated as described in *A*. The data are shown as the mean value ± SEM. *N* = 2 independent experiments. Number of analyzed cells (*n*) = 78 (ctrl), 54 (shTSC1), 58 (shTSC1 +p70S6Ki). \*\*\**p* < 0.001, \*\**p* < 0.01 (Kruskal Wallis test [H=100.9] with Dunn’s multiple comparisons test). **C.** Representative confocal images of CCD-1070Sk transduced with control (pLKO) or shTSC1-encoding lentiviral vector in basal conditions (not stimulated with PDGF), immunofluorescently stained for PDGFR (magenta) and EEA1 (green). Scale bar = 10 µm.

**Figure S2.**
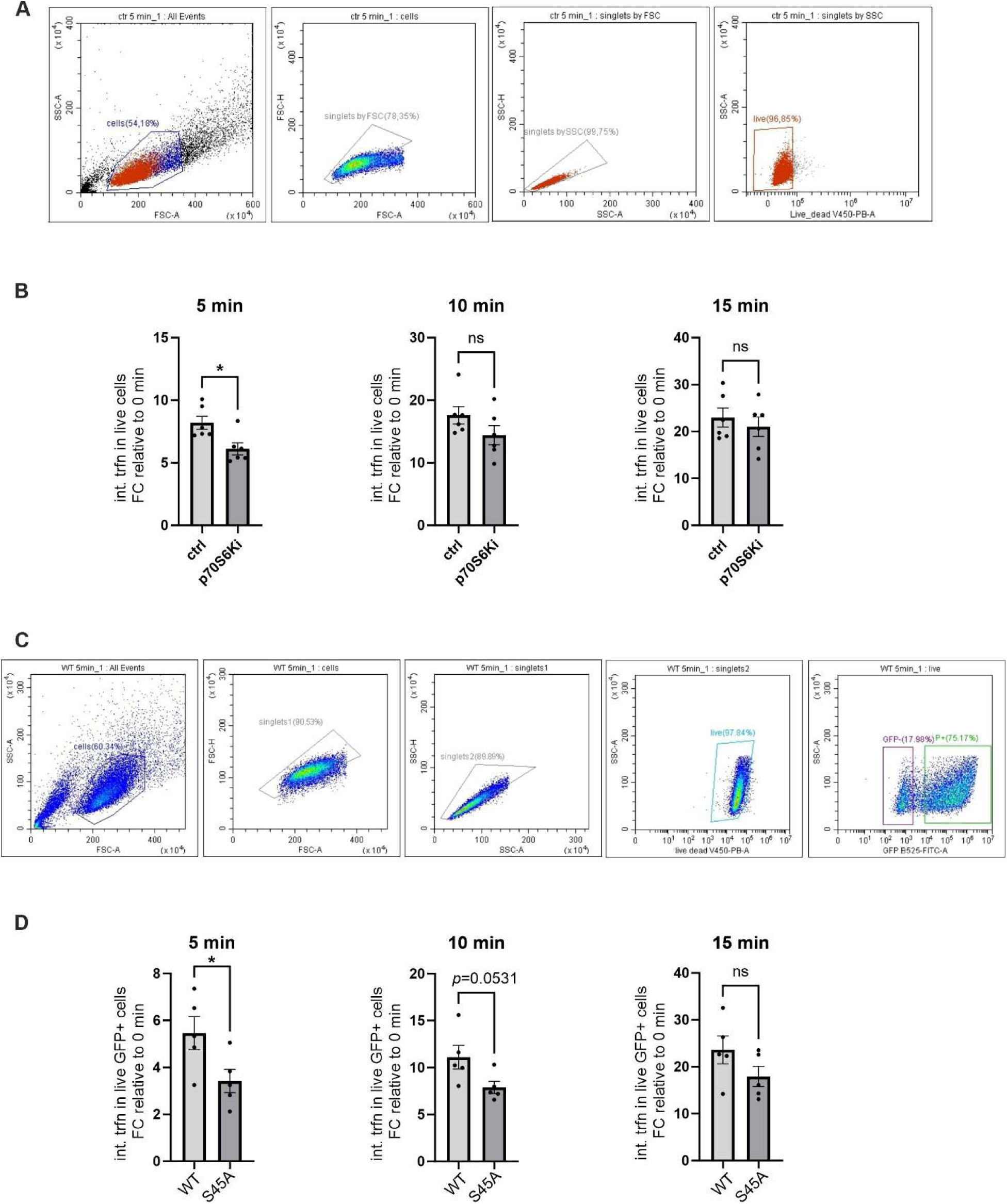
Evaluation of transferrin uptake by flow cytometry. **A.** Representative example of gating strategy for transferrin uptake measurement in HeLa cells treated with DMSO (ctrl) or p70S6K inhibitor (p70S6Ki). **B.** Quantification of the median fluorescence intensity of internalized Alexa Fluor 647-conjugated transferrin after 5, 10, and 15 min in cells treated as in panel *A*. Data are presented as mean values normalized to the 0 min time point ± SEM. *N =* 6 independent experiments. Unpaired t-test (5 min: *\*p* < 0,05 t = 2,950, df = 10; 10 min: *ns*-non-significant, t = 1,554, df = 10; 15 min: *ns* – non-significant t = 0,6692, df = 10). **C.** Representative example of gating strategy for transferrin uptake measurement in HeLa KO µ2 cells transfected with plasmids encoding EGFP-μ2^WT^ or EGFP-μ2^S45A^ **D.** Quantification of the median fluorescence intensity of internalized Alexa Fluor 647-conjugated transferrin after 5, 10, and 15 min in cells treated as in panel *C*. Data are presented as mean values normalized to the 0 min time point ± SEM. *N* = 5 independent experiments. Unpaired t-test (5 min: *\*p* < 0.05; t = 2.368, df = 8; 10 min: *ns –* non-significant, t = 2.267, df = 8; 15 min: *ns –* non-significant, t = 1.546, df = 8).

**Figure S3.**
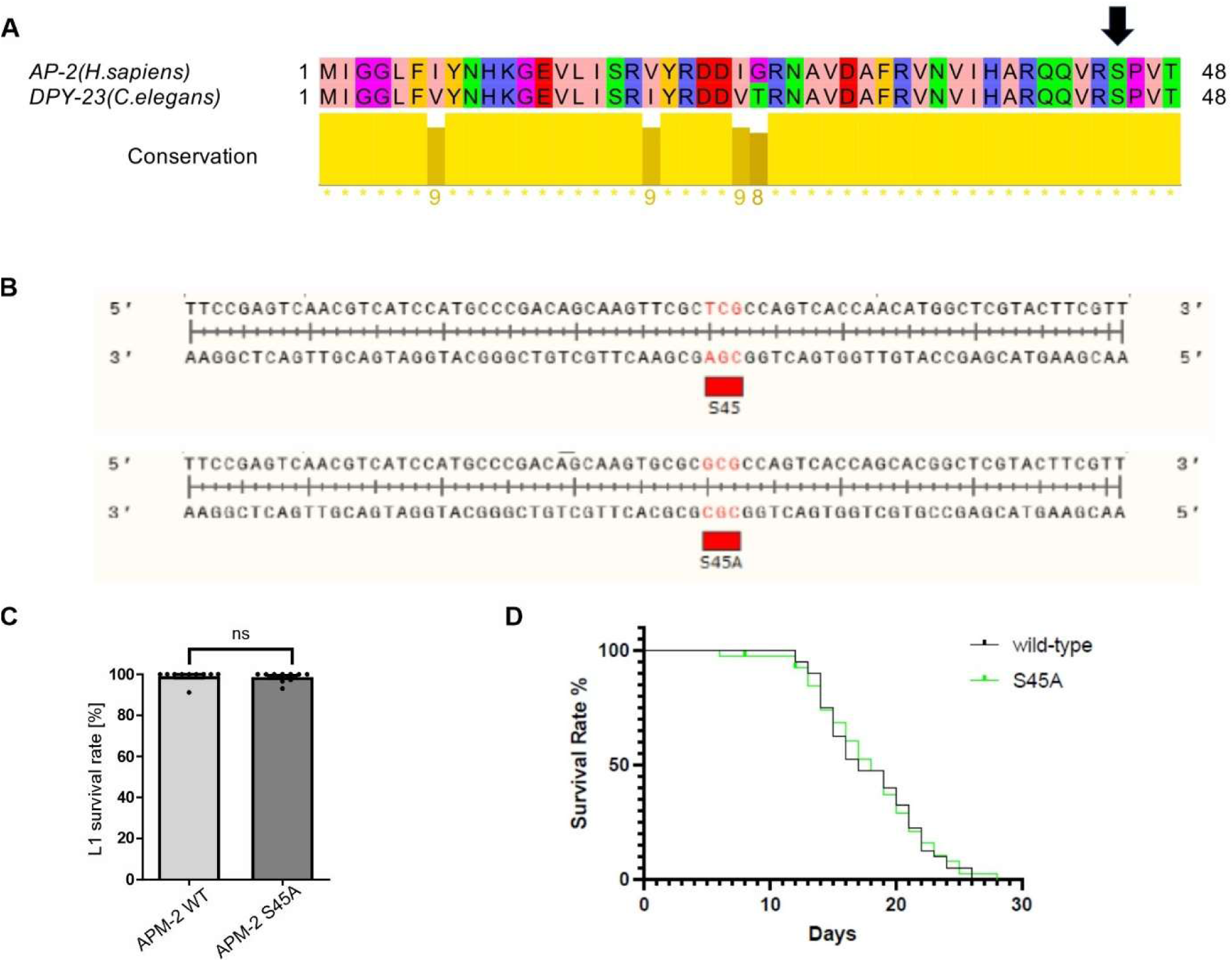
Serine 45 in APM-2 in *C. elegans.* **A.** Global alignment of N-terminus of human AP2 (UniProt ID: Q96CW1) and *C. elegans* DPY-23 (UniProt ID: P35603). Sequences were aligned using the EMBOSS Needle web server (Madeira *et al*, 2022) with default parameters and visualized in the Jalview Desktop software (Waterhouse *et al*, 2009) with residues colored by their physicochemical properties; the arrow indicates S45 position. **B.** Representative sequencing results for the R160.1b.1 locus in the wild-type (N2) strain (top section) compared with nematodes where the 375th position was edited to change T to C, resulting in the amino acid substitution to Ala45 (A). The data were examined using SnapGene 6 software (Insightful Science; snapgene.com). **C.** Percentage of hatched L1 larvae from eggs laid over 24 hours by wild type (N2) and apm-2 mutant strain. *ns* – non-significant (Mann–Whitney test). **D.** Lifespan comparisons of control (WT – wild type, N2) and APM-2S45A worms at 20°C. *ns* – non-significant (Mantel-Cox log-rank test).

**Figure S4.**
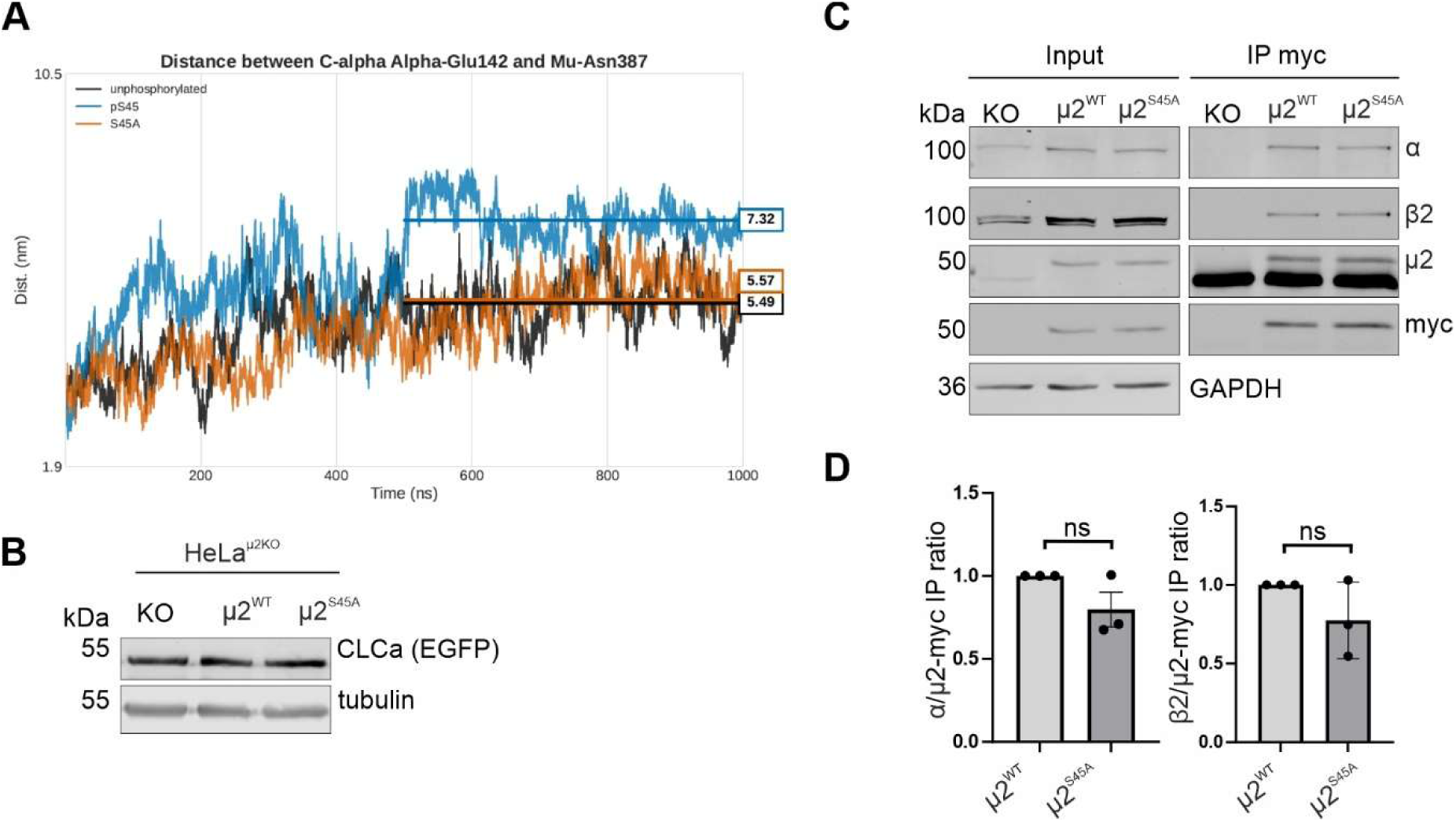
The impact of the µ2 S45A mutation on the AP2 complex and the level of CLCa. **A.** Effects of μ2 S45 phosphorylation and S45A mutation on AP2 core in open conformation. Time evolution of the distance between the C-alpha atoms of α-Glu142 and μ2-Asn387, a metric used to monitor the opening of the AP2 core, over 1000 ns all-atom MD simulations. Traces represent the unphosphorylated (black), P-S45 (blue) and S45A (orange) states. **B.** Western blot showing level of EGFP-fused clathrin light chain (CLCa) and tubulin in HeLa KO μ2 cells stably expressing EGFP-CLCa and myc-µ2^WT^ or myc-µ2^S45A^. **C.** Western blot showing co-immunoprecipitation of myc-μ2 with other AP2 subunits: α and β2 from HeLa KO μ2 cells stably expressing EGFP-CLCa and myc-µ2^WT^ or myc-µ2^S45A^. Input, 10% of lysate used for immunoprecipitation. Shown is a representative example of *N =* 3 independent experiments. **D.** Quantitative analysis of immunoprecipitated AP2 α and β2 subunits. *N* = 3 independent experiments. *ns* – non-significant (one sample t-test; α [t=1,940, df=2], β2 [t=1,602, df=2]).

**Figure S5.**
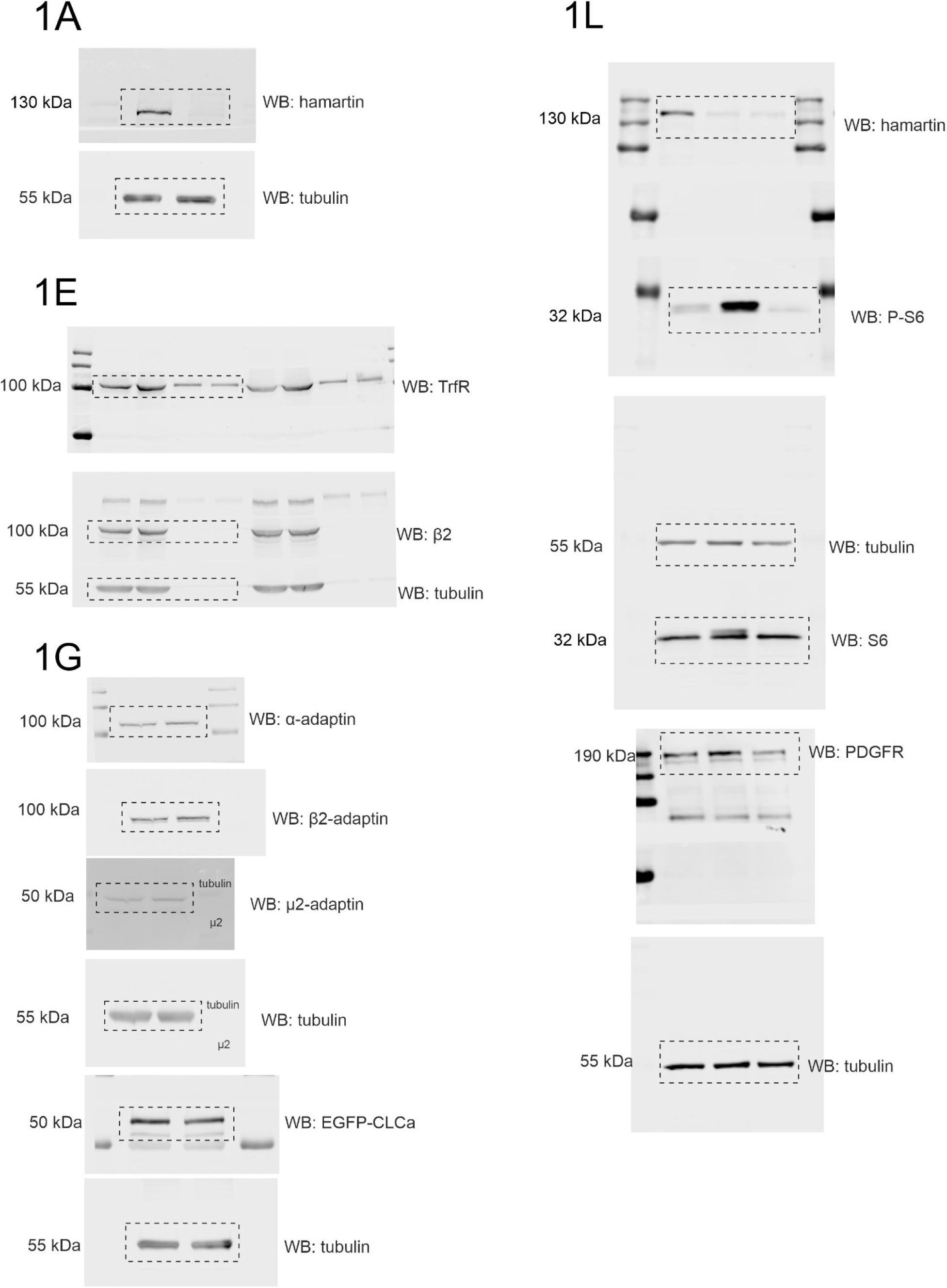
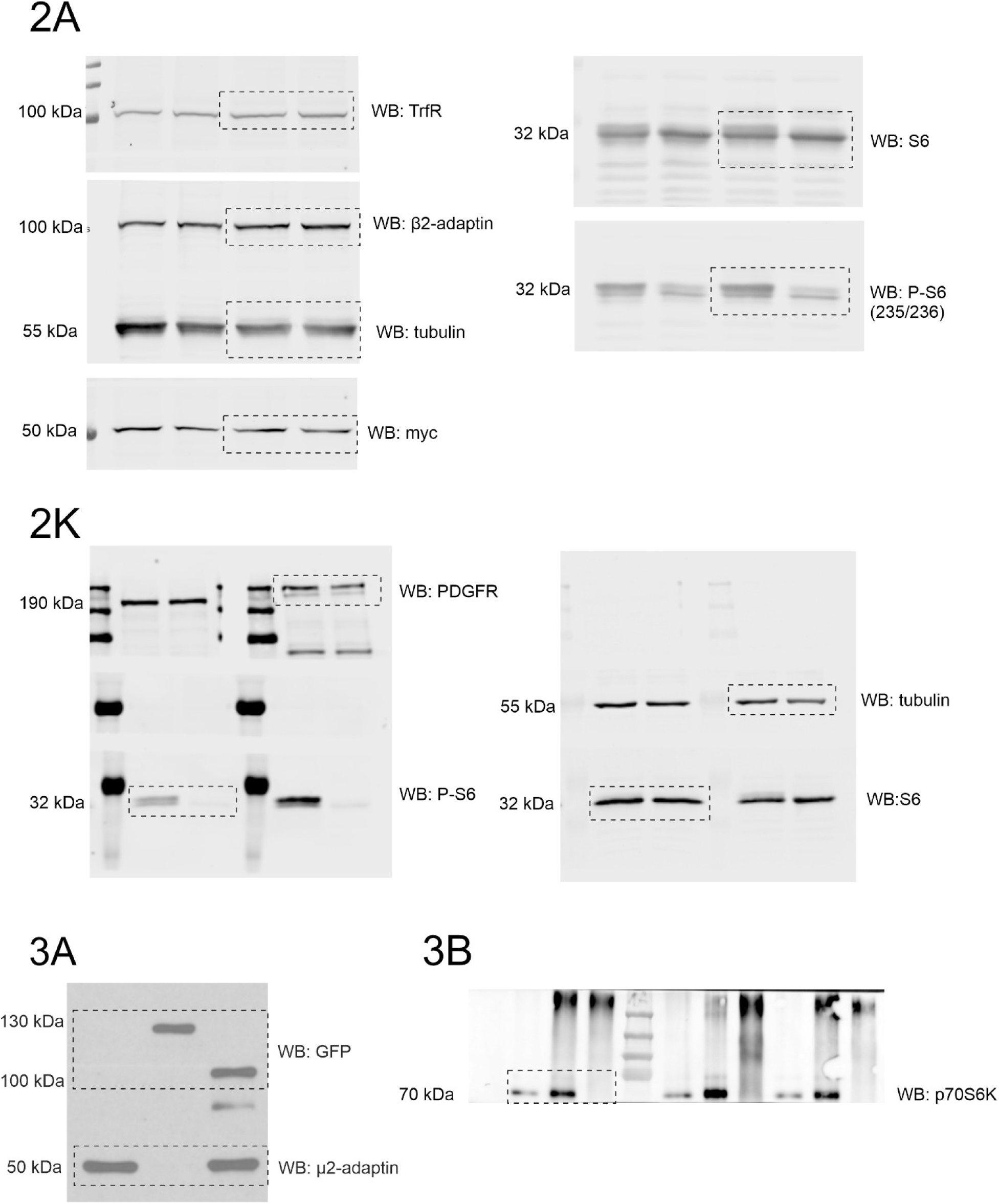
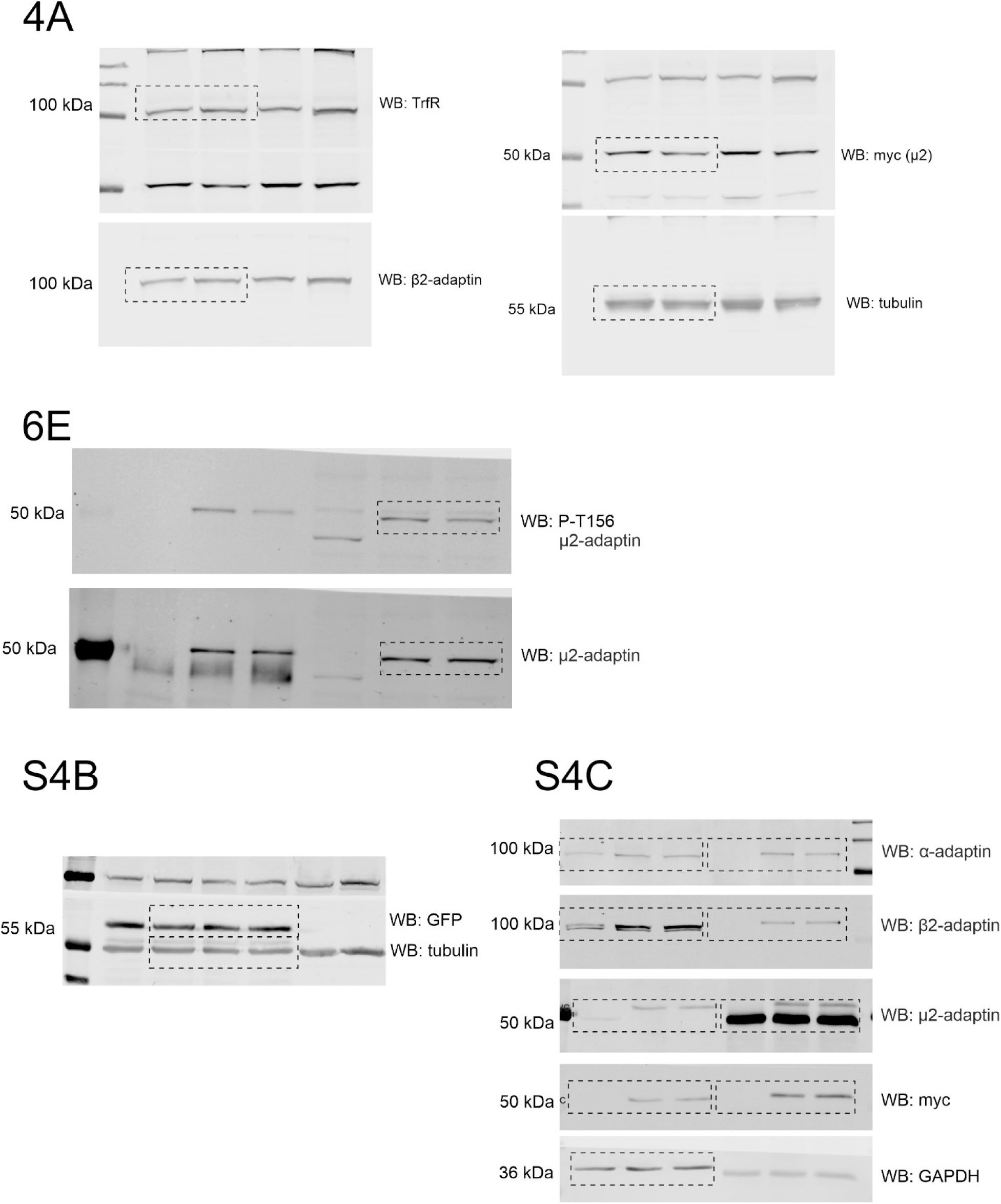
Blot transparency. Full-size blots that correspond to cropped images that are presented in the manuscript.

**Table S1.**
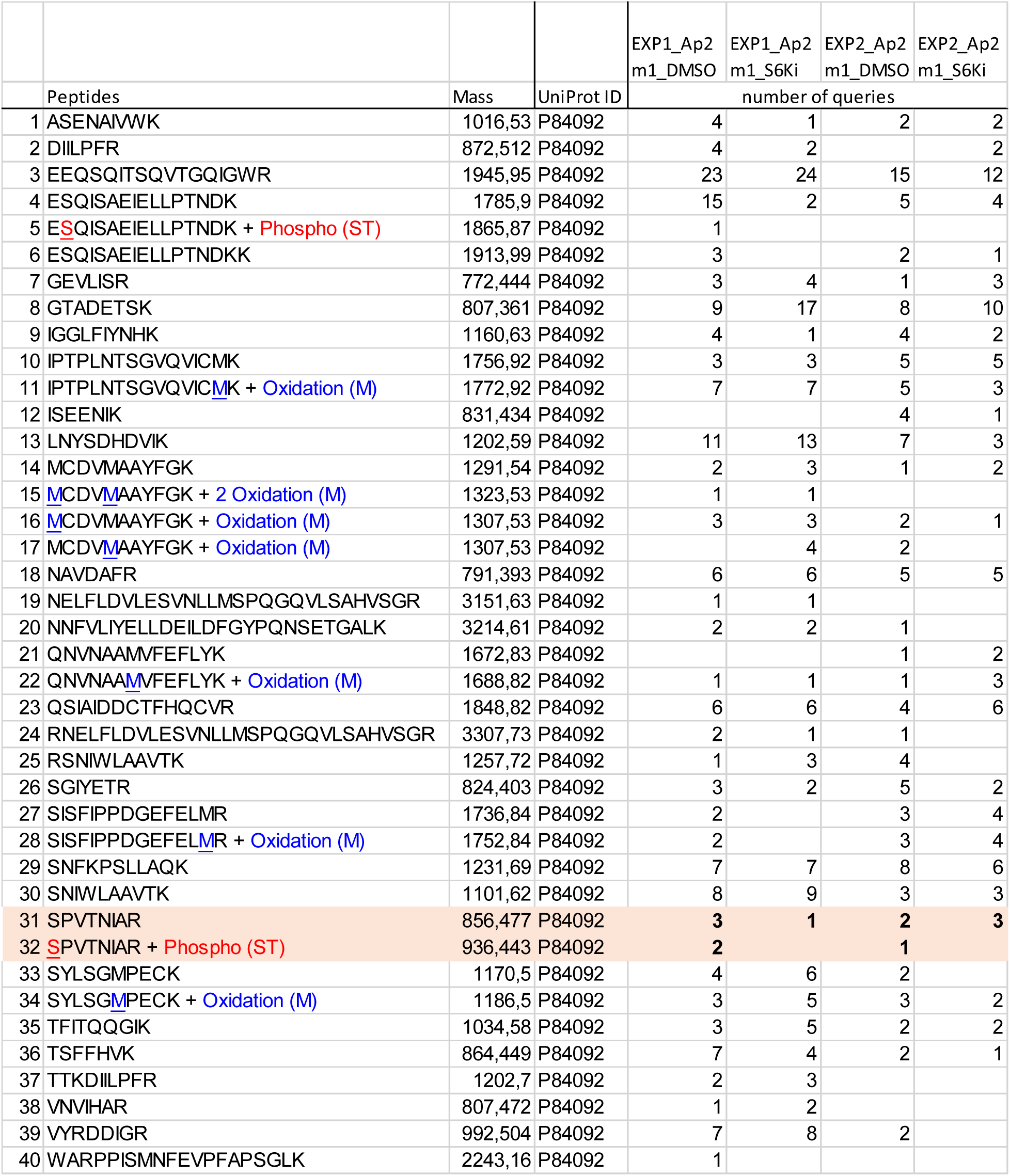
Comparison of detected μ2 peptides and post-translational modifications between DMSO and p70S6Ki-treated samples.

## Notes

### Competing Interest Statement

The authors have declared no competing interest.

### Summary of Updates

New data on transferrin uptake evaluated by flow cytometry were added.

